# *Trypanosoma brucei* ribonuclease H2A is an essential enzyme that resolves R-loops associated with transcription initiation and antigenic variation

**DOI:** 10.1101/541300

**Authors:** Emma Briggs, Kathryn Crouch, Leandro Lemgruber, Graham Hamilton, Craig Lapsley, Richard McCulloch

## Abstract

In every cell ribonucleotides represent a threat to the stability and transmission of the DNA genome. Two types of Ribonuclease H (RNase H) tackle such ribonucleotides, either by excision when they form part of the DNA strand, or by hydrolysing RNA when it base-pairs with DNA, in structures termed R-loops. Loss of either RNase H is lethal in mammals, whereas yeast can prosper in the absence of both enzymes. Removal of RNase H1 is tolerated by the parasite *Trypanosoma brucei* but no work has examined the function of RNase H2. Here we show that loss of the catalytic subunit of *T. brucei* RNase H2 (TbRH2A) leads to growth and cell cycle arrest that is concomitant with accumulation of nuclear damage at sites of RNA polymerase (Pol) II transcription initiation, revealing a novel and critical role for RNase H2. In addition, differential gene expression of both RNA Pol I and II transcribed genes occurs after TbRH2A loss, including patterns that may relate to cytosolic DNA accumulation in humans with autoimmune disease. Finally, we show that TbRH2A loss causes R-loop and DNA damage accumulation in telomeric RNA Pol I transcription sites, leading to altered variant surface glycoprotein expression. Thus, we demonstrate a separation of function between the two nuclear *T. brucei* RNase H enzymes during RNA Pol II transcription, but overlap in function during RNA Pol I-mediated gene expression during host immune evasion.

---

Incorporation of ribonucleotides is a major threat to the stability of DNA genomes. Such incorporation can occur in three ways, each of which can be tackled by ribonuclease H (RNase H) enzymes. Ribonucleotide monophosphates (rNMPs) can be directly incorporated into DNA by DNA polymerases (Pols), an error that occurs at various frequencies depending on the selectivity of dNTPs over rNTPs by the different types of DNA Pol and by the base type [1–3]. The ratio of rNTPs/dNTPs also influences rNMP selection, with rNTPs exceeding dNTPs in the cellular pool [4]. These factors result in as many as 13,000 and 3 million rNMPs being incorporated into the yeast and human genomes per round of replication, respectively [1, 5, 6]. Once incorporated, ribonucleotides destabilise DNA due to the presence of a reactive 2’-hydroxyl group on the ribose sugar, rendering the DNA backbone more vulnerable to cleavage. rNMPs are further incorporated as RNA primers necessary for the initiation steps of DNA replication. Whereas leading strand replication initiates from a single origin and a single RNA primer, lagging strand replication requires 7-14 nucleotide RNA primers for the synthesis of each Okazaki fragment, which is normally ∼200 bp in length [7–9], meaning ribonucleotides are found through this DNA strand. RNA is also frequently found associated with genomic DNA in the form of R-loops: for instance, nascent RNA can become hybridised to the template DNA strand behind a transcribing RNA Pol, forming a heteroduplex and displacing a single strand of DNA [10]. Alternatively, R-loops can form in *trans* when RNA generated at one genomic location forms an RNA-DNA hybrid elsewhere in the genome [11–13]. Though R-loops have been linked to several genomic functions[10, 14, 15], including transcription, DNA replication, chromosome segregation and telomere homeostasis, the RNA-DNA hybrids can also lead to instability and mutation [16–18], particularly when RNA biogenesis is compromised [19–23] and at sites of clashes between the DNA replication and transcription machineries, potentially contributing to replication fork collapse [24, 25].

All organisms encode RNase H enzymes that degrade RNA incorporated in DNA [26]. Though RNase H enzymes can contribute to the removal of DNA replication-associated RNA primers, two other nucleases, flap endonuclease 1 (FEN1) and Dna2, appear to play a larger role in ensuring these ribonucleotides remain only transiently in DNA [7, 27–29]. In contrast, RNase H enzymes play a more critical role in removing embedded ribonucleotides and R-loops. Most organisms encode two RNase H enzymes, type 1 and type 2. Eukaryotic type 2 RNase H is termed RNase H2 and is a complex made up of catalytic subunit, A, and two further subunits, B and C. In contrast, RNase H1, the type 1 enzyme, is a monomer. Only RNase H2 is able to remove embedded ribonucleotides, which it does by initiation of the ribonucleotide excision repair (RER) pathway [30, 31]. In this reaction, RNase H2 detects the 2’-OH group and cleaves 5’ of an embedded ribonucleotide, resulting in a DNA nick. DNA Pol δ subsequently performs PCNA-dependent nick translation and displaces the ribonucleotide, which is then removed by FEN1. Finally, DNA ligase repairs the lesion. In contrast to the specific role of RNase H2 in RER, both eukaryotic RNase H enzymes are able to resolve R-loops [32], which they do by hydrolysing the RNA within the RNA-DNA hybrid. In yeast, both the RER and R-loop activities of RNase H2 are known to protect against genomic instability [33, 34], although the protein is not essential for cell viability, even when gene mutation is combined with loss of RNase H1 [34]. In contrast, RNase H1 and RNase H2 are both essential for mouse embryonic development: lack of the former impairs mitochondrial DNA replication, while lack of the latter results in increased levels of ribonucleotides and DNA lesions in the nuclear genome. In addition, in humans, mutations in all three RNase H2 subunits have been shown to cause the auto-inflammatory disease Aicardi– Goutières syndrome (AGS) [35]. Mice lacking fully functional RNase H2 display features of AGS that result from aberrant activation of an innate immune pathway that normally targets foreign cytosolic DNA [36, 37]. However, what self-molecules cause aberrant autoimmunity in these AGS models, how they reach the cytosol, and what activity of RNase H2 causes their generation, remains unclear [38].

In previous work we examined the distribution of R-loops across the genome of *T. brucei*, comparing mammal-infective wild type (WT) cells with mutants lacking *T. brucei* RNase H1 (TbRh1). The *T. brucei* genome, in common with all kinetoplastids [39], is arranged radically differently from most eukaryotes, since virtually all protein-coding genes (∼8,000) are expressed by RNA Pol II from a relatively small number of multigenic transcription units, meaning each gene does not have its own defined promoter, but many (sometimes hundreds) of genes share a transcription start site, where conserved RNA Pol II promoter sequences have not been found. Genes within a single polycistronic transcription unit (PTU) are initially encoded as a potentially multigene pre-mRNA before mature mRNAs are generated by coupled 5’ RNA *trans*-splicing (adding the cap) and polyadenylation [40]. RNA Pol II transcription initiation, as well as termination, has been mapped to so called strand switch regions (SSRs), which separate adjacent PTUs, including by RNA-seq [41] and chromatin immunoprecipitation (ChIP) of modified and variant histones [42], modified base J [43], and a subunit of RNA Pol II [44]. Transcription in *T. brucei* also appears functionally linked with DNA replication, since at least one component of the origin recognition complex (ORC) binds to SSRs, although only a subset are activated during S phase [45, 46]. We have shown that the most abundant site of R-loop accumulation in the constitutively transcribed *T. brucei* ‘core’ genome is at intergenic sequences within the RNA pol II PTUs, where the RNA-DNA hybrids display precise association with regions of low nucleosome density, suggesting a relationship with polyadenylation and, perhaps, *trans*-splicing [47]. R-loops are also notably enriched at SSR boundaries where RNA Pol II transcription initiates, whereas little or no R-loop signal is observed at SSRs where transcription termination takes place. Although R-loops increase at various loci after deletion of TbRH1 [47], no cell cycle or growth defects are observed, even though it might be predicted that R-loops in the PTUs present an obstacle to replication and transcription, or that they might form at potentially predictable sites of clashes between the *T. brucei* replication and transcription machineries, perhaps suggesting TbRH1-independent mechanisms to avoid such conflict.

Gene expression in *T. brucei* displays further novelty in that some proteins are transcribed by RNA Pol I, not II. Whilst resident in the mammalian host, trypanosomes express a dense ‘coat’ of variant surface protein (VSG), from one of ∼15 telomeric multigene RNA Pol I VSG bloodstream form expression sites (ES) [48]. Co-transcribed with the VSG are multiple expression site-associated genes (ESAGs), most of which also encode surface proteins [49]. In order to evade host immunity *T. brucei* continually switches between expression of antigenically distinct VSGs, a process termed antigenic variation [50]. One mechanism for VSG switching is silencing transcription from the single active ES and activating transcription from a previously silent ES, containing a different VSG. Additionally, recombination mechanisms allow VSG sequences (from ∼2,000 genes and pseudogenes) to be copied from silent subtelomeric arrays, minichromosomes or the silent ES into the active ES [50, 51]. R-loops are found at low levels in the active ES of WT *T. brucei*, suggesting they form co-transcriptionally[52]. Upon deletion of *TbRH1*, R-loop signal significantly increases in both active and silent ES, and is associated with the accumulation of DNA damage in the active ES and increased levels of expression of previously silent VSGs [52]. Though these data implicate R-loops targeted by TbRH1 in VSG switching, how the RNA-DNA hybrids are linked to transcriptional and recombinational switching, and whether they are solely recognised by TbRH1 is unknown. For instance, increased DNA damage is mainly detected in S and G2 phase TbRH1 mutant parasites, but if and how DNA replication converts R-loops into ES DNA breaks is unknown [46, 53]. Moreover, overexpression of TbRH1 has been shown to decrease VSG switching in TbRAP1-depleted cells [54], but how such a telomere function relates to VSG transcription and recombination is unclear.

Here we describe the function of RNase H2 in *T. brucei*, which we examined to attempt to clarify how R-loop formation and resolution contributes to RNA Pol II core transcription and to VSG transcription and recombination. We show that loss of the *T. brucei* RNase H2A catalytic subunit, TbRH2A, is lethal and leads to cell cycle stalling associated with extensive nuclear DNA damage, but without loss of DNA synthesis. Mapping reveals that DNA damage accumulates specifically at transcription initiation sites after loss of TbRH2A, which also causes a decrease in R-loops at the same loci. Loss of TbRH2A also causes R-loop and DNA damage accumulation across the VSG ES, with increased changes in VSG expression. Finally, RNA-seq details differential gene expression of both RNA Pol I and II transcribed genes after TbRH2A loss. Thus, we demonstrate a separation of function between the two nuclear *T. brucei* RNase H enzymes in the context of multigene RNA Pol II transcription, but overlap in functions during antigenic variation.

## Results

### RNase H2A is an essential nuclear protein in bloodstream form *T. brucei*

In order to identify putative type 2 RNase H proteins in *T. brucei*, BLAST and protein domain analyses were employed, searching the *T. brucei* genome with both type 1 and type 2 RNase H proteins from *E. coli* and a range of eukaryotes (Table S1). As we [52] and others [55] have described, a single RNase H1 can be readily detected in *T. brucei* (TbRH1). In addition, three candidates for the *T. brucei* RNase H2 complex were revealed: Tb427.10.5070 was predicted to encode a protein highly similar to eukaryotic catalytic RNase H2A subunits, and Tb427.01.4220 and Tb427.01.4730 encode likely orthologues of RH2B and RH2C, respectively (Table S1, Fig.S1A). Surprisingly, synteny between these orthologues and three previously described RNase H2-like genes in *Leishmania major* [56] is not simple to discern. For instance, the putative *T. brucei* RNase H2A (TbRH2A) encoded by Tb427.10.5070 by shows greatest homology to, and is syntenic with, LmjF.36.0640, which has previously been named RNase HIIB [56](Fig. S1B). Moreover, Tb427.10.5070 is not syntenic with LmjF.13.0050, despite this gene being predicted to encode a type 2 RNase H (named RNase HIIA)[56]. The predicted amino acid sequence of Tb427.10.5070 revealed conservation of active site and catalytic residues described in type 2 RNase H enzymes in other organisms (Fig. S1B), consistent with the gene encoding the catalytic subunit of RNase H2. To begin to test for function, TbRH2A was C-terminally tagged with 6 copies of the HA epitope and expressed from its endogenous locus in mammalian-infective (bloodstream form; BSF) parasites (Fig.S2B, C). Immunofluorescence with anti-HA antiserum revealed signal throughout the nucleus in all cell cycle stages and without any discernible sub-nuclear localisation (Fig.S2C), features shared with TbRH1 [52], suggesting the presence of two RNase H enzymes in the *T. brucei* nucleus. Also in common with TbRH1, TbRH2A-6HA fluorescence signal increased in cells undergoing nuclear DNA replication (Fig. S2D).

Attempts to generate *TbRH2A* null mutants were unsuccessful (data not shown), suggesting the protein is essential. To test this prediction, we employed tetracycline (tet)-inducible RNA interference (RNAi) [57]. *TbRH2A* transcript levels were reduced to ∼3% of the parental cells when TbRH2A^RNAi^ cells were cultured in the presence of tet for 36 hr (Fig.1A). Indeed, *TbRH2A* RNA levels were ∼19% lower in TbRH2A^RNAi^ cells than in the parental cells even when grown in the absence of tet, indicating some expression of the RNAi-inducing *TbRH2A* stem-loop RNA prior to induction, consistent with some altered phenotypes prior to RNAi induction (described below). Nonetheless, tet-induction of RNAi caused a severe growth defect relative to uninduced cells, with cell proliferation stalling 24 hr post-induction (Fig.1B). Cell cycle progression (as analysed by staining nuclear (n) and kinetoplast (k) DNA with DAPI) was also severely altered: after 24 hr of RNAi induction, the proportion of cells with one nucleus and two kinetoplasts (1N2K) increased to ∼40% of the population (Fig.1C), representing a ∼4-fold increase compared with uninduced cells (where ∼10% were 1N2K) and suggesting a stall in nuclear G2/M phase. Small numbers of cells with 1 nucleus and more than 2 kinetoplasts (1NXK) could also be detected at 24 hrs, and increased substantially at 30 hr induction, indicating kDNA replication continued after depletion of TbRH2A (Fig.1C, D). Similarly, the small numbers of cells (∼3%) seen after 24 hr induction that had multiple nuclei or aberrant nuclear staining, as well as >2 kinetoplasts (YNXK), increased after 30 hrs of RNAi (Fig.1C, D). These complex perturbations in growth after loss of TbRH2A, suggestive of a partial cell cycle stall and some further, ineffective nuclear replication and division, appeared not to increase from 36-72 hrs (Fig.1C), consistent with the lack of population growth or death during this time (Fig.1B). To further investigate the effects of TbRH2A depletion on the cell cycle, flow cytometry was used to examine DNA content (Fig.1E, F). Reduction in cells with 2N content (G1 phase) was apparent, consistent with the loss of 1N1K cells in the DAPI staining. However, the proportion of cells with 4N content (G2/M) decreased until 36 hr of tet-induction (∼18.2%), which is inconsistent with the increased number of 1N2K cells seen by DAPI. An explanation most likely lies in the pronounced increase of cells detected with more than diploid genome content (>4N, from ∼7% at 0 hr to ∼47 % at 36 hr), suggesting that many of the cells scored as 1N2K, and apparently stalled in the cell cycle at G2/M, continued to synthesise nuclear DNA but mainly failed to effectively execute mitosis.

**Figure 1.**
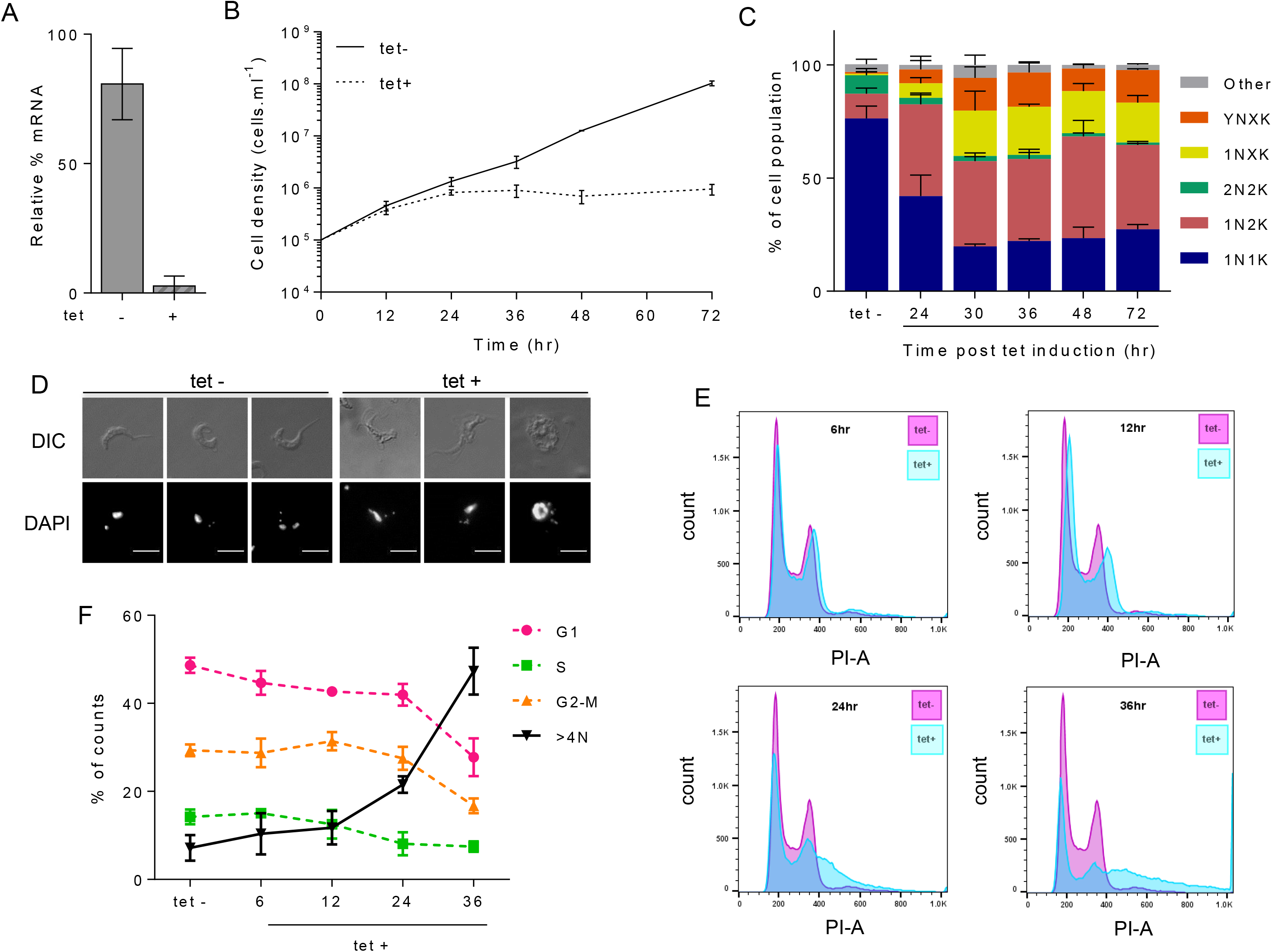
RNase H2A is essential for bloodstream form *T. brucei* viability. A) Levels of *TbRH2A* transcripts in tetracycline induced (+) and uninduced (-) cells after 24 hr of culture, relative to 2T1 cells (levels set at 100%), determined by RT-qPCR; error bars show SD of two independent experiments. B) Cumulative growth curves of tet+ and tet-TbRH2A RNAi cultures, showing cell densities over 72 hr. C) Bar graph showing, at multiple time points, the percentage of tet-induced cells in the population that correspond to the following cell types, defined by DAPI staining of the nucleus (N) and kinetoplast (K): 1N1K, 1N2K, 2N2K, 1NxK (>2 K foci), YNXK (>2 K foci and aberrant N number or morphology), and other (do not conform to the above). Tet – shows the average of uninduced samples from all time-points. D) Example images of (tet +) induced and un-induced (tet -) cells after 30 hr of growth; scale bar, 5 μm. E) Profiles of propidium iodide (PI) stained uninduced (tet-, pink) and RNAi induced (tet +, blue) populations after 6, 12, 24 and 36 hr growth; y-axes show cell counts, and x-axes shows PI-area fluorescence. F) Graph showing the percentage of cells in each expected cell cycle stage (G1, S and G2-M), or cells with genome content >4N, based on measuring proportion of the population with 2N, 2N-4N, 4N and >4N content; tet-shows the average of all tet - time points. In B), C) and F), error bars shown SD of three independent experiments.

### DNA synthesis continues in RNase H2A depleted *T. brucei* despite increased nuclear DNA damage

To ask if loss of TbRH2 affects nuclear genome functions, we measured levels of DNA damage before and after TbRH2 RNAi using antiserum recognising Thr130-phosphorylated histone H2A (γH2A), a known marker of DNA damage in *T. brucei* [58, 59]. Consistent with previous reports [58], immunofluorescence (IF) with anti-γH2A antiserum revealed nuclear signal in a small fraction (∼10%) of uninduced cells (Fig.2A,B). In contrast, IF of cells grown for 12, 24 and 36 hr in the presence of tet (Fig.2A,B) revealed dramatic time-dependent increases in nuclear γH2A signal, reaching ∼75% of the RNAi induced cells in the population after 36 hr (Fig.2B). Western blotting of whole cell protein extracts confirmed the IF data, with γH2A levels showing large increases 24 and 36 hrs after RNAi induction compared with uninduced cells (Fig. 2C). To explore how such widespread nuclear damage relates to replication of the nuclear genome, we incubated RNAi induced and uninduced parasites with the thymine analogue 5-ethynyl-2′-deoxyuridine (EdU) and detected its incorporation via Click-IT chemistry (Fig.2D,E; Fig.S3). ∼99% of cells, cultured in the absence of tet, incorporated EdU in the nucleus after a 4 hr incubation with the analogue (Fig.2E, S3), which is as expected for asynchronous BSF *T. brucei* cells with a cell cycle time of ∼6 hr, since virtually all cells should, at least partially, undergo nuclear S phase and take up EdU. Even after 36 hr of RNAi induction, when population growth had stopped (Fig.1B), ∼93% of cells incorporated EdU (Fig.2D,E; Fig.S3A). Hence, loss of TbRH2A had little, if any effect, on DNA synthesis (Fig.1E,F). Moreover, since virtually all the RNAi-induced cells that had γH2A signal in their nucleus had also incorporated EdU, with overlap of the two signals (Fig.2D; Fig.S3A), the extensive nuclear DNA damage caused by loss of TbRH2 did not appear to impede DNA replication, consistent with flow cytometry indicating increased DNA content in the absence of effective mitosis (Fig.1C,E,F).

**Figure 2.**
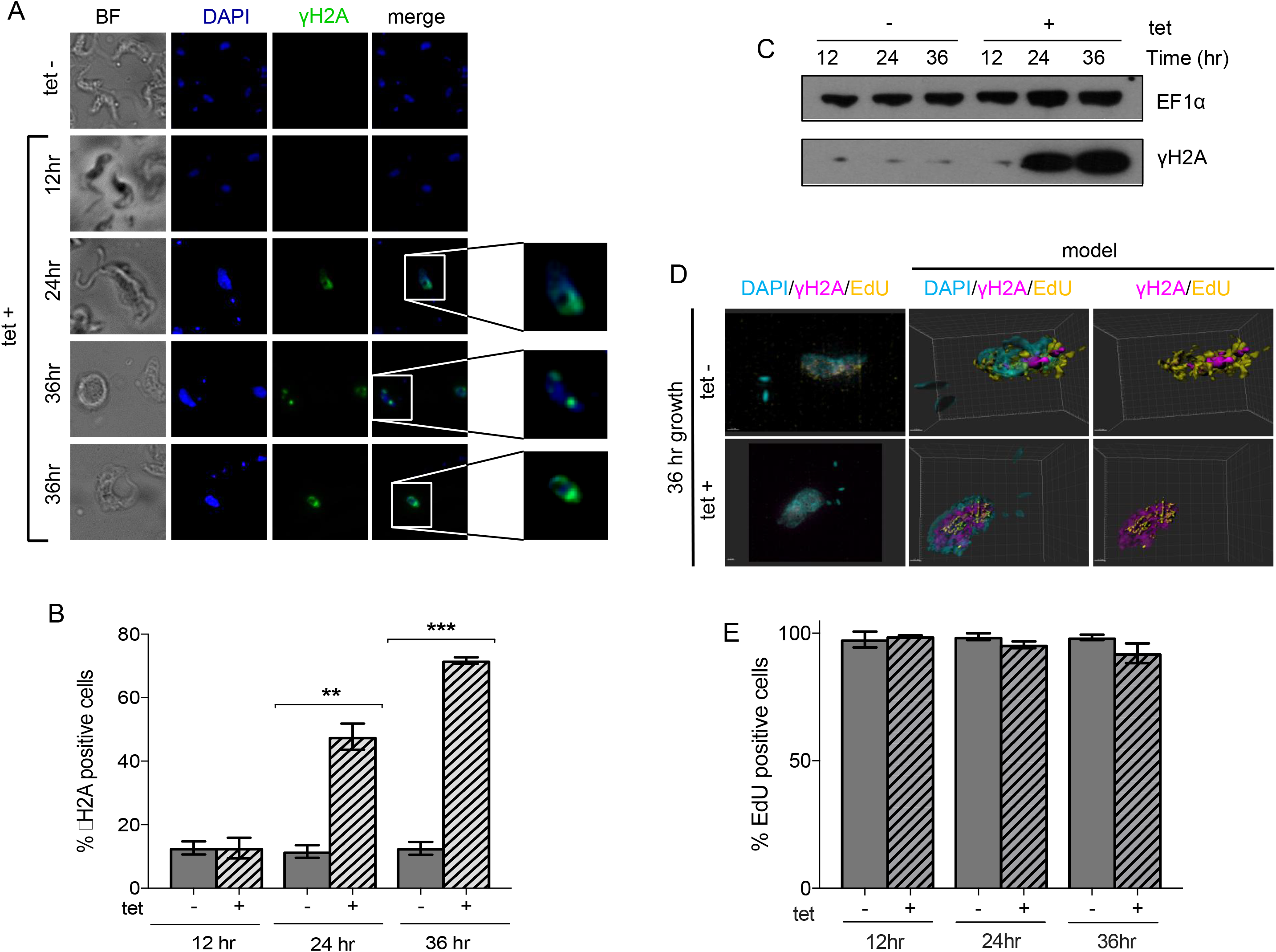
TbRH2A-depleted parasites accumulate DNA damage yet continue to synthesise DNA. A) High-resolution imaging of DAPI (blue) and γH2A (green) immunofluorescence with (tet +) and without (tet -) RNAi induction, at various time points; images to the right show increased magnification of the boxed nuclear DNA. B) Bar graphs showing the percentage of tet + and tet - populations positively staining for γH2A after 12, 24 and 36 hrs growth; error bars show SD of three independent experiments. C) Western blot detection of γH2A in whole cell protein extracts after 12, 24 and 36 hrs growth of tet + and tet - cells; EF1α staining is shown as a loading control. D) Example SR-SIM images of DAPI, γH2A and EdU staining are shown, along with 3D reconstructions (model). TbRH2A RNAi cells were cultured in the presence (tet +) or absence of tet (tet -) for 36 hr before imaging; scale bar = 1 μm. E) Bar graph showing the percentage of tet + and tet - populations positively staining for EdU incorporation after 12, 24 and 36 hrs of growth; error bars show the SD for three independent experiments (further examples of immunofluorescence images of EdU and γH2A staining are shown in Fig. S3).

### R-loop abundance at transcription start sites decreases after depletion of RNase H2A

To ask if the extensive nuclear genome damage seen after TbRH2A RNAi relates to R-loop distribution, we used monoclonal antibody S9.6 [60] to immunoprecipitate DNA-RNA hybrids from formaldehyde-fixed chromatin derived from RNAi-induced or uninduced cells, evaluating distribution by mapping reads to the *T. brucei* genome after next generation DNA sequencing (DRIP-seq). Fig.S4 shows genome-wide DRIP-seq mapping after 24 hr of growth with and without tet-induced RNAi, revealing widespread R-loop enrichment and some correlation with DNA repeats. To understand how R-loop distribution in the TbRH2A^RNAi^ cells compared with similar analysis in *TbRH1* null mutants and WT *T. brucei* cells [47], DRIP-enriched regions were defined as locations with ≥1.2 fold-change increase in IP mapped reads relative to pre-IP samples. Analysis of these enriched regions showed they were mainly found in RNA Pol II PTUs (∼88% of uninduced, and ∼86% of RNAi induced; Fig.S5), a very similar distribution to that previously reported for WT and *TbRH1* null mutant DRIP-seq data. Moreover, DRIP enriched regions within the PTUs, before and after TbRH2A RNAi, were most clearly associated with intergenic sequences (∼58% of uninduced, and ∼60% of RNAi induced samples; Fig.S6A,B), a localisation bias that appeared slightly increased compared with WT cells (∼50% of intra-PTU enriched regions; Fig.S6A). Correspondingly, the number of DRIP enriched regions associated with gene coding DNA sequence (CDS) was reduced in both the TbRH2A uninduced (6,715 regions) and, even more so, in the RNAi induced (5,300 regions) DRIP-seq data compared with WT (8861 regions; Fig.S6A). Heatmaps of DRIP-seq enrichment around every RNA Pol II gene confirmed the predominant enrichment around the CDS, with relatively precise signal localisation upstream and downstream of each CDS, as was seen in WT cells (Fig.S7) and *TbRH1* null mutants [47]. Taken together, these data indicate relatively stable accumulation of R-loops within the RNA Pol II PTUs, which is not markedly altered by loss of TbRH2A or TbRH1 [47]. Indeed, outside the RNA Pol II PTUs, DRIP-seq of both induced and uninduced TbRH2A^RNAi^ cells revealed R-loop enrichment in Pol I and Pol III transcribed genes, retrotransposon hotspot (RHS) genes, and in centromeres (Fig.S4, S5), in each case at comparable levels to WT cells and TbRH1 null mutants [47], suggesting many of the R-loops that form in the *T. brucei* genome are relatively unaffected by loss of either RNase H activity. However, within this context of global R-loop stability, two regions displayed notable changes in DRIP-seq profile after TbRH2A loss: transcription start sites, and VSG genes (as described below).

Previously, we described pronounced accumulation of DRIP-seq signal around the sites of transcription initiation in the SSRs that separate adjacent RNA Pol II PTUs [47]. Here, mapping of DRIP-seq in the same loci revealed differences in signal between the tet-treated RNAi-induced cells and both RNAi uninduced and WT cells. Fig.3A shows DRIP-seq signal plotted over every SSR, with the loci separated into the following classes: divergent SSRs, representing sites of transcription initiation in both sense and antisense directions; convergent SSRs, which are sites of transcription termination; and head-to-to tail SSRs, where transcription both terminates and initiates on the same strand. Fig.3B provides detailed mapping at examples of each class of SSR. In all cases DRIP-seq signal was largely equivalent between the uninduced and WT cells. Unexpectedly, upon depletion of TbRH2A, DRIP-seq signal was substantially reduced compared with uninduced and WT cells around the two locations of transcription initiation in divergent SSRs, and at the single site of transcription initiation in the head-to-tail SSRs (right and central panels, Fig.3A,B). In contrast, the same loss of DRIP-seq signal was not detected at the convergent SSRs, or at the locations of transcription termination in head-to-tail SSRs (central and left panels, Fig.3A,B). To examine this effect of TbRH2A loss change further, genes were separated into those predicted to be first within a PTU (i.e. proximal to the transcription start sites; n, 110) and all others (n, 8278; internal to the PTU) and the pattern of DRIP-seq examined around the genes’ ATG (Fig.S8). For all genes DRIP-seq abundance peaked upstream of the ATG and was depleted downstream of the ATG. Moreover, like we described in *TbRH1* null mutants [47], increased DRIP-seq was seen upstream of the ATG in the uninduced TbRH2A^RNAi^ cells compared with WT, perhaps due to ‘leaky’ RNAi causing some loss of TbRH2A. In contrast, after TbRH2A RNAi a notable loss of DRIP-seq signal was seen around the ATG of transcription start site-proximal genes but not PTU-internal genes, consistent with loss of the RNase H mainly affecting R-loop abundance around sites of transcription initiation. Next, we re-grouped the SSRs according to whether or not they have been identified as origins of DNA replication [45] and re-analysed DRIP-seq distribution (Fig.S9). This analysis revealed reduction of DRIP-seq signal across both types of SSRs in the RNAi induced cells compared with uninduced and WT, indicating that it is transcription, and not DNA replication, of the SSRs that dictates the change in DRIP-seq signal after loss of TbRH2A.

**Figure 3.**
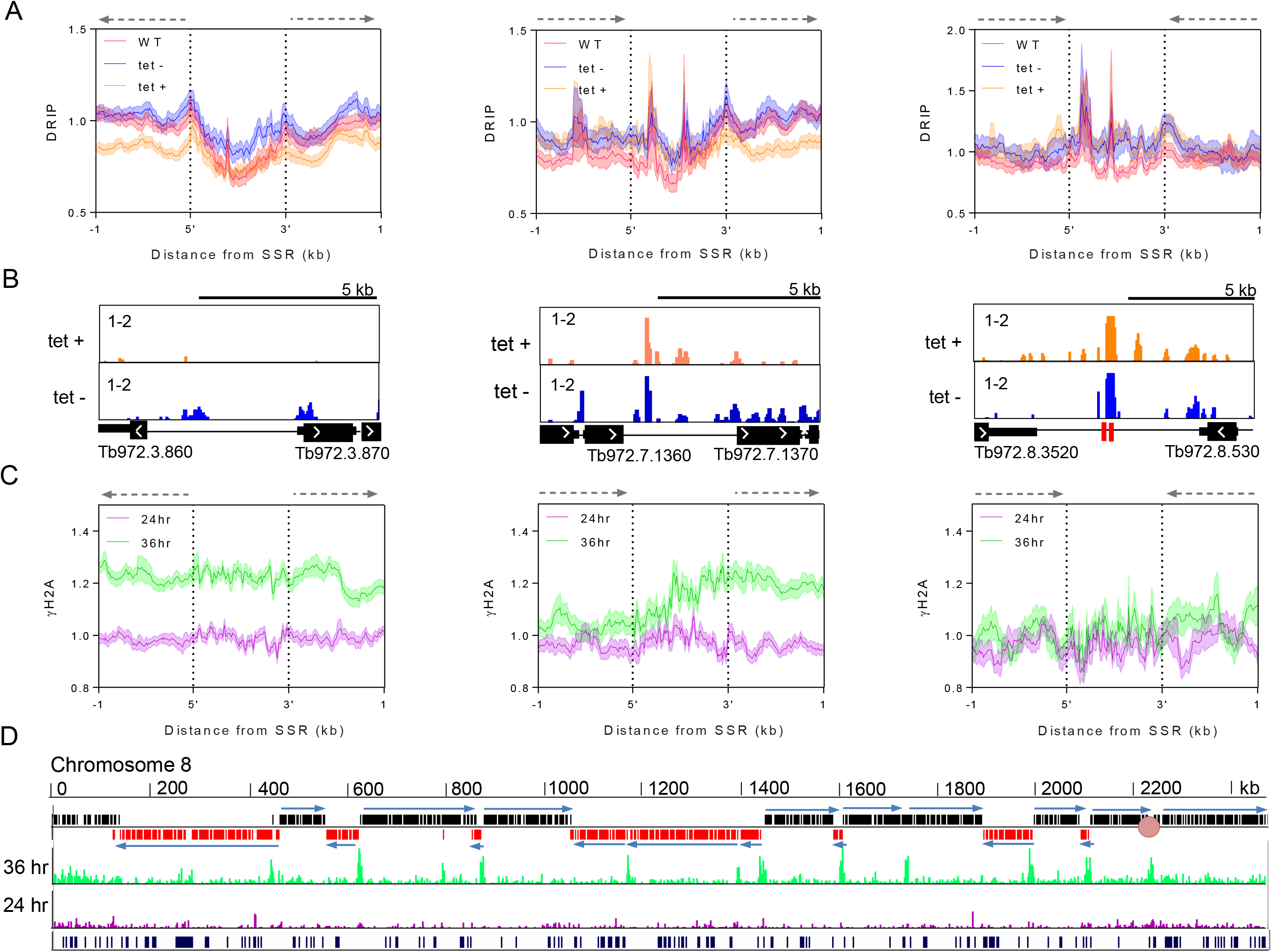
After TbRH2A depletion RNA-DNA hybrids decrease, and DNA damage increases at sites of RNA Pol II transcription initiation. A) Average DRIP-seq signal is shown as metaplots plotted for WT (pink), TbRH2A RNAi un-induced (tet -, blue) and RNAi induced (tet +, orange) data sets over divergent (left), head-to-tail (middle) and convergent (right) SSRs (+/- 1 kb). In all cases 5’ and 3’ denote SSR boundaries defined by flanking transcript coordinates. Transcription direction is shown above the plots by dashed arrows. Standard error is shown as shaded regions. B) Example screenshots of DRIP-seq signal in tet + and tet - cells at individual SSRs in each class; CDS (thick black), UTR (thin black lines) and snRNA/tRNA genes (red) are shown below the DRIP-seq tracks. C) Metaplots of γH2A ChIP-seq signal in TbRH2A RNAi induced samples relative to un-induced is shown after 24 hr (pink) and 36 hr (green) of RNAi induction; average signal is plotted across SSRs as for (A). D) γH2A ChIP-seq signal in induced relative to un-induced cells is also shown plotted across chromosome 8 after 24 hr (pink) and 36 hr (green) of growth (scale 1-3 fold-change). Upper track shows genes on sense (black) and antisense (red) strands, and arrows highlight transcription direction; the lowest track shows tandem repeat sequences.

### TbRH2A RNAi induces DNA damage at transcription start sites in the core genome and results in gene expression changes

Given the highly localised changes in R-loop abundance after TbRH2A RNAi and the pronounced accumulation of γH2A, we next sought to ask if the effects are connected. To address this, we performed chromatin-immunoprecipitation (ChIP)-seq with anti-γH2A antiserum to map sites of accumulation of the modified histone, comparing read depth in tet-induced and uninduced cells grown for 24 and 36 hrs (Fig.3C,D; Fig.S10). In all cases read depth in the IP samples was first normalised with pre-IP (input) samples, before fold-change in the RNA-induced cells was calculated relative to uninduced at each time point; Fig. S10 shows the resulting ratios of γH2A ChIP signal across the whole genome. Little change in γH2A localisation or signal was detected 24 hrs after TbRH2A RNAi induction (Fig.3C, D; Fig.S10), despite increased γH2A signal in IF and western analyses (Fig.2). However, after 36 hr of RNAi induction, clear increases in γH2A signal could be discerned that coincided with the boundaries of the PTUs in the core genome (Fig.3D). To ask if this accumulation was specific for transcription start sites, the γH2A ChIP-seq data was mapped to SSRs grouped, as before, into divergent, convergent and head-to-tail classes (Fig.3C). Increased reads were clearly apparent 36 hrs after RNAi across the convergent SSRs and around the sites of transcription initiation in the head-to-tail SSRs, but increased reads could not be detected at convergent SSRs. Hence, increase in the DNA damage marker closely correlates with the locations of RNA-DNA hybrid loss, albeit with R-loop decrease seen before the accumulation of Thr130-phosphorylated H2A. Nonetheless, these data indicate that depletion of TbRH2A has localised effects, not seen after ablation of TbRH1 [47], which connect R-loops and DNA damage at sites of multigenic transcription initiation in *T brucei*.

To ask if TbRH2A loss after RNAi affects gene expression, RNA-seq analysis was conducted, comparing mRNA abundance after 24 and 36 hrs of RNAi relative to uninduced control cells. After 24 hrs of TbRH2A RNAi, remarkably few changes in gene expression were seen: no genes displayed significantly reduced RNA abundance, while 32 showed significantly increased abundance (Fig.4A,B). More marked changes were found 36 hr post-induction: 113 gene-specific RNAs significantly increased in abundance, and 396 were significantly reduced (Fig.4A,B). Interestingly, and as described further below, in keeping with our previous finding that R-loops can induce VSG switching in *T. brucei* [52], 30 of the up-regulated genes after 24 hr of TbRH2A depletion were annotated as VSGs (5) or ESAGs (25), a number that increased further after 36 hrs (37 VSGs, 14 ESAGs). At the same time more procyclin genes were significantly increased, suggesting effects on RNA Pol I transcription (Fig. 4B), as well as increased numbers of RHS-associated genes, where R-loops could be mapped (Fig.1).

**Figure 4.**
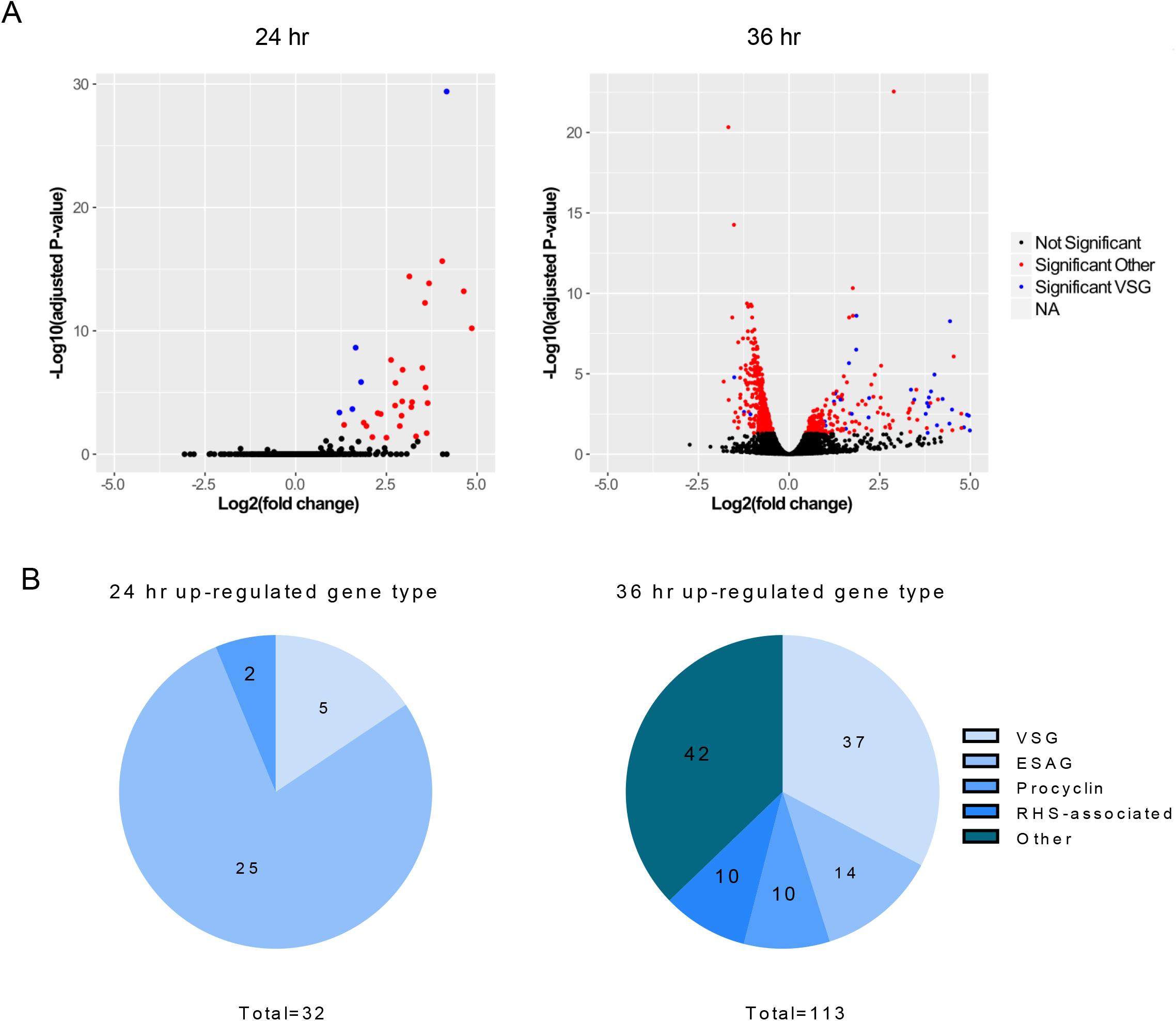
Gene expression is altered upon depletion of TbRH2A. A) Volcano plots displaying differential expression of genes after TbRH2A knockdown. Each data point represents a gene. Genes were deemed significantly differentially expressed when RNA-seq indicated an adjusted p value < 0.05 of gene-specific RNA in induced cells relative to uninduced. X-axes show the log2 fold-change between un-induced and induced after 24 hr (left) and 36 hr (right) of culture, and y-axes shows log2 adjusted p value. Data was generated with two independent replicates for each condition and time point. Significantly differentially expressed genes are shown in red or blue (denoting VSG); all other genes, including VSGs, are shown in black. B) Number of genes significantly up-regulated in tet-induced TbRH2A RNAi cells relative to uninduced, after 24 (left) and 36 hr (right) of growth, are shown annotated as VSGs, ESAGs, procyclin, RHS-associated, and other genes; total numbers are shown below each chart.

The remaining 42 up-regulated genes detected after 36 hrs RNAi, as well as the larger number of down-regulated genes, were more diverse in function (Fig.S11). However, GO term analysis revealed that several of the down-regulated genes were involved in small molecule biosynthesis pathways, and prominently represented were genes involved in nucleotide and ribonucleotide synthesis and salvage (Fig. S11). Other terms found to be over-represented in the down-regulated gene set were metabolic processing of other small molecules, including cellular carbohydrates, ketones and organic acids (Fig. S11). We could find no correlation between altered RNA abundance and gene position within a PTU.

### R-loops and DNA damage accumulate in VSG expression sites after RNase H2A depletion

We have previously reported increased BES-associated R-loops in *TbRH1* null mutant BSF *T. brucei*, coinciding with increased levels of VSG switching and increased replication-associated DNA damage [52]. To ask if these antigenic variation-directed effects are limited to TbRH1 activities, we first plotted DRIP-seq before and after TbRH2A RNAi across all the available BES sequence [48], comparing the mapping to DRIP-seq in WT cells (Fig.5A,B; Fig.S12). To limit cross-mapping of short reads to related BES we applied MapQ filtering [61]. After 24 hr, both uninduced and induced cells showed substantially increased DRIP-seq signal across both the active (BES1) and all inactive BESs (Fig.5A,B; Fig.S12) in comparison with WT. Increased signal in the uninduced cells is presumably due to leaky expression of the TbRH2A-targetting stem-loop RNA, consistent with the slight reduction in TbRH2A RNA (Fig.1A). Strikingly, the pattern of R-loop distribution in the BES compared well with TbRH1 null mutants, with prominent signal in the 70 bp-repeats and little localisation to sequences between the ESAGs, distinguishing the RNA-DNA hybrid distribution from the RNA Pol II PTUs (Fig.6A, Fig.S12). Surprisingly, we found no evidence for further increases in DRIP-seq signal after RNAi induction, for reasons that are unclear. To establish if the DRIP-seq detects RNA-DNA hybrids, and to ask if the structures increase in abundance after prolonged RNAi depletion of TbRH2A, DRIP-qPCR was performed with samples prepared after 36 hr of growth with and without tet induction (Fig.5B), including, in each case, with a parallel IP that was treated with *E. coli* RNase HI (EcRH1) to degrade the RNA within the hybrids. ESAGs 6 and 8 were targeted to examine R-loops within all BES, while primers recognising VSG221 and VSG121 were used to examine the predominantly transcriptionally active (BES1) BES and one transcriptionally silent BES (BES3), respectively. In all cases, increased RNA-DNA hybrids were detected in the RNAi induced samples relative to uninduced, indicating DRIP-seq may underestimate the effect of TbRH2A loss in the BES, and treatment with EcRH1 reduced IP levels for nearly all samples, validating detection of R-loops (Fig. 5B).

**Figure 5.**
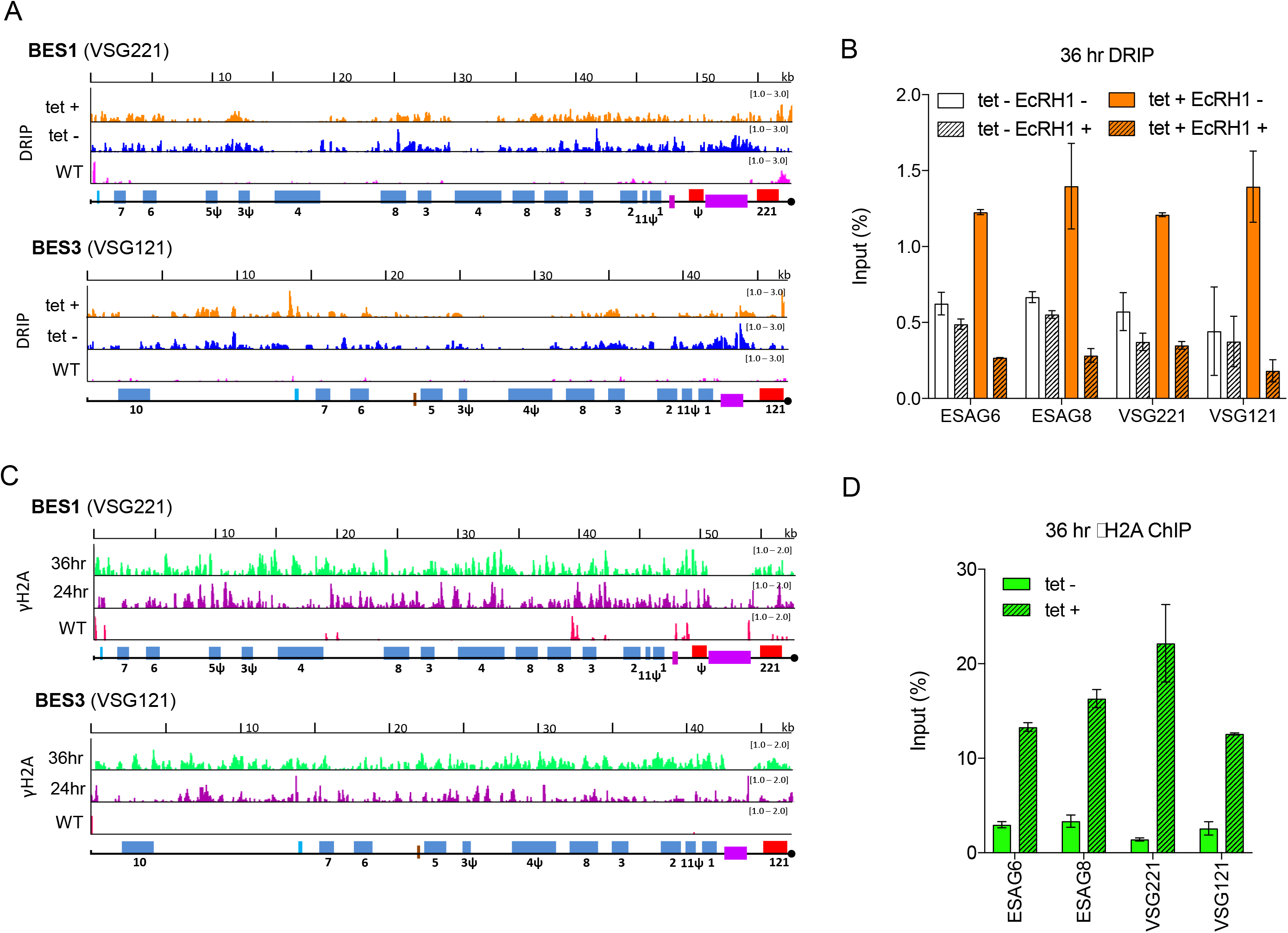
R-loop and yH2A levels increase across VSG BESs in cells depleted of TbRH2A. A) Localisation of R-loops is shown in BES1 (active site of VSG transcription in WT cells) and BES3 (one normally transcriptionally silent site), comparing DRIP-seq signal in TbRH2A RNAi cells grown for for 24hr in the absence (tet -; blue) or presence (tet +; orange) of RNAi induction, and compared with WT cells (pink). BES features are shown as follows: promoter (aqua), ESAGs (blue, numbered), 70-bp repeats (purple) and VSGs (red); pseudogenes are indicated (Ψ), and the end of the available BES sequence is denoted by a black circle. B) DRIP-qPCR using primers targeting the sequences of ESAG6, ESAG8, VSG221 (BES1) and VSG121 (BES3), with or without *E.coli* RNase HI (EcRHI) treatment, showing the percentage of amplification in the IP sample relative to input from tet induced (tet+) and non-induced (tet -) cells grown for 36 hr. Error bars display SEM for three replicates. C) yH2A ChIP-seq signal enrichment is shown mapped to BES1 and BES3 as a ratio of reads in tet-induced samples relative to un-induced (each first normalised to the cognate input sample) after 24 (purple) and 36 (green) hr growth; γH2A ChIP-seq signal (normalised to input) is shown in WT cells for comparison (pink). D) γH2A ChIP-qPCR, as in (B): data is shown for tet induced (tet +) and non-induced (tet -) cells after 36 hr of growth. Error bars display SD for two replicates.

To ask if the BES R-loops also correlate with damage in this component of the genome, we next mapped γH2A ChIP-seq to the BESs, comparing signal fold-change in the RNAi-induced samples relative to uninduced (Fig.5C,D; Fig.S13). These data revealed a number of features. First, extensive accumulation of γH2A signal was seen across the entire length of both active and inactive BESs (Fig.5C), as confirmed by DRIP-qPCR (Fig.5D). The extent of the accumulation, and the presence of phosphorylated H2A in both active and inactive BES, is distinct from the γH2A ChIP-seq profile seen in *TbRH1* null mutant cells [52], where γH2A is only significantly mapped to telomere-proximal regions of the active BES, not the silent sites. Second, accumulation of γH2A in the BES was clearly discernible 24 hrs post RNAi induction, unlike in the core genome, where signal was only seen after 36 hrs RNAi. In addition, accumulation of the modified histone was not limited to, or more pronounced at, the promoter of the BES. Thus, loss of TbRH2A has a more rapid and widespread effect on BES integrity than is seen in the core genome. Third, in contrast with the strong DRIP-seq signal across the 70 bp repeats after TbRH2A loss, γH2A ChIP-seq was notably limited on this BES feature relative to all other parts of the BESs, perhaps indicating a particular effect of repeat composition on generation or spreading of the modified histone.

### Loss of RNase H2A leads to changes in the VSG expression

As discussed above, RNA-seq analysis revealed increased RNA levels of several VSG, ESAGs and procyclin genes in response to TbRH2A depletion (Fig. 4). To examine this in more detail, we first used RT-qPCR to examine levels of VSG221, which is expressed from the predominantly active BES, and four VSGs housed in normally transcriptionally silent BESs, after 24 and 36 hr of TbRH2A RNAi (Fig. 6A). Levels of VSG221 transcript did not change significantly after TbRH2A knockdown, but levels of two silent VSGs increased after 24 hrs RNAi, and all four silent VSG levels increased relative to uninduced samples after 36 hr of TbRH2A knockdown (Fig. 6A). To investigate changes in VSG transcription more widely, RNA-seq reads were mapped to all the BESs (Fig. 6C, Fig.S14), as well as a collection of 2470 VSG sequences described in the *T. brucei* Lister 427 genome [62](Fig.6B,D), and differential expression analysis was repeated. 20 VSG sequences were found to be significantly up-regulated after 24 hr of TbRH2A depletion, a cohort that increased to 50 VSGs after 36 hr of RNAi (Fig. 6B). Of these, 40% and 60% were classified as pseudogenes in the 24 hr and 36 hr samples, respectively. Of the remaining VSG sequences up-regulated after 24 hr of RNAi, 25% were classified as intact genes found within subtelomeric arrays, 20% as intact genes associated with mini-chromosomes, 10% as metacyclic (M) ES VSGs, and the remaining 1 VSG as BES-housed (Fig. 6B). After 36 hr, a similar proportion of up-regulated VSGs were array-associated (26%), 10% were housed in the BESs, and 2% (1 VSG) were MES-associated (Fig. 6B); no mini-chromosome associated VSGs were significantly up-regulated at this time point. Hence, VSG sequences from across the diverse genome repertoire were found to be transcribed after TbRH2A depletion. Within the silent BESs, RNA-seq mapping suggested that not only did VSG RNA levels increase after RNAi, but also levels of promoter-proximal ESAGs (most clearly seen as ESAG 6 and 7; Fig.6C), explaining the increased levels ESAG-associated reads described in Fig. 4.

**Figure 6.**
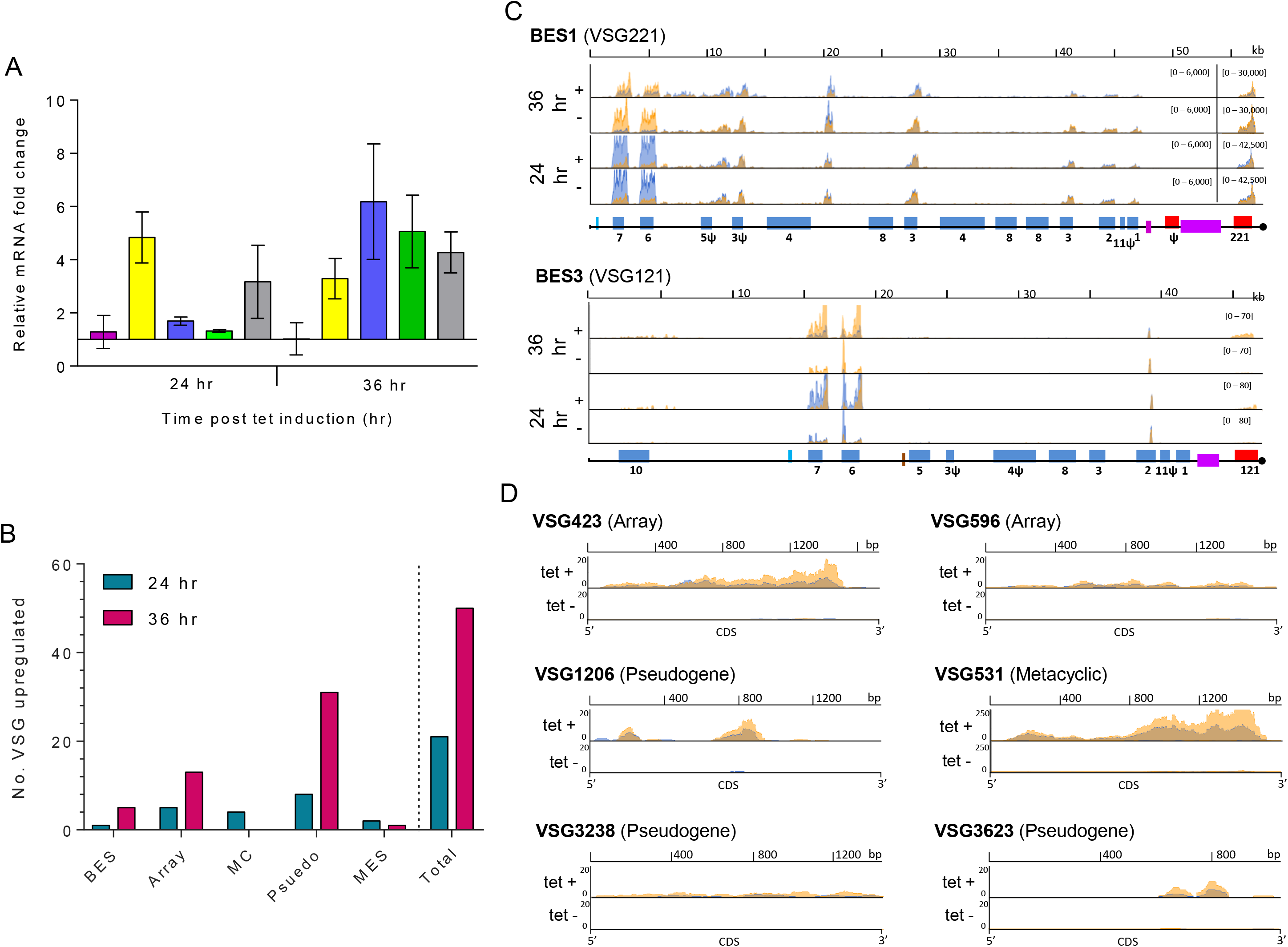
Depletion of TbRH2A results in increased transcription of silent VSGs. A) Graph of RNA levels for 5 VSGs, determined by RT-qPCR, in tet-induced TbRH2A RNAi cells, plotted as fold-change relative to levels of the cognate VSG RNA in un-induced cells after both 24 hr and 36 hr of culture; VSG221 (pink) is in the active BES (BES1) of WT cells, while VSG121 (yellow), VSG800 (blue), VSGT3 (green) and VSG13 (grey) are in silent BESs; error bars show SD for three independent experiments. B) Graph depicting the number of VSG genes whose RNA levels display significant upregulation in RNAi-induced RNA-seq samples relative to un-induced, both 24 hr and 36 hrs after growth; the total number is sub-categorised depending on whether the VSGs have been localised to the BES, are intact genes in the subtelomeric arrays (array), are in minichromosomes (MC), are pseudogenes (pseudo), or are in a metacyclic VSG ES (MES). C) Normalised RNA-seq read depth abundance (y-axes) is plotted for two independent replicates (overlaid orange and blue) in TbRH2A RNAi parasites after 24 and 36 hr of growth, with (tet +) or without (tet -) RNAi induction. Read depth is shown relative to gene position (x-axes) for BES1 and BES3. D) As in (C), showing for RNA-seq read depth abundance (y-axes) across a selection of non-BES VSG CDS (x-axes), after 36 hr of growth.

In order to ask if changes in VSG RNA levels in response to TbRH2A depletion extended to VSG protein changes on the parasite surface, expression of two VSGs, VSG221 (active BES1) and VSG121 (inactive BES3), was analysed via immunofluorescence with specific antisera (Fig.7A,B). Unpermeabilised cells were probed for expression of both VSGs after 12, 24 and 36 hr growth with and without tet-induction of RNAi. In the absence of RNAi, across all time points, virtually all parasites solely expressed VSG221 (∼99%) (Fig. 7B), with only 1% detected that did not stain for either VSG. After 12 hr of induction, singular VSG expression did not significantly change. However, after 24 hr of RNAi nearly 2% of cells did not stain for either VSG, and ∼0.2% expressed both VSGs simultaneously. By 36 hr of induction, the number of cells expressing both VSG221 and VSG121 increased further to ∼0.7% of the population, parasites expressing neither VSG also increased to ∼4.5% and, at this time point alone, a small number of cells (∼0.2%) could be detected that expressed VSG121 but not VSG221. Taken together, these data indicate that loss of TbRH2A results in an increased frequency at which *T. brucei* cells inactivate expression of VSG221, as well as causing loss of the controls that ensure monoallelic expression of a single VSG in one cell.

**Figure 7.**
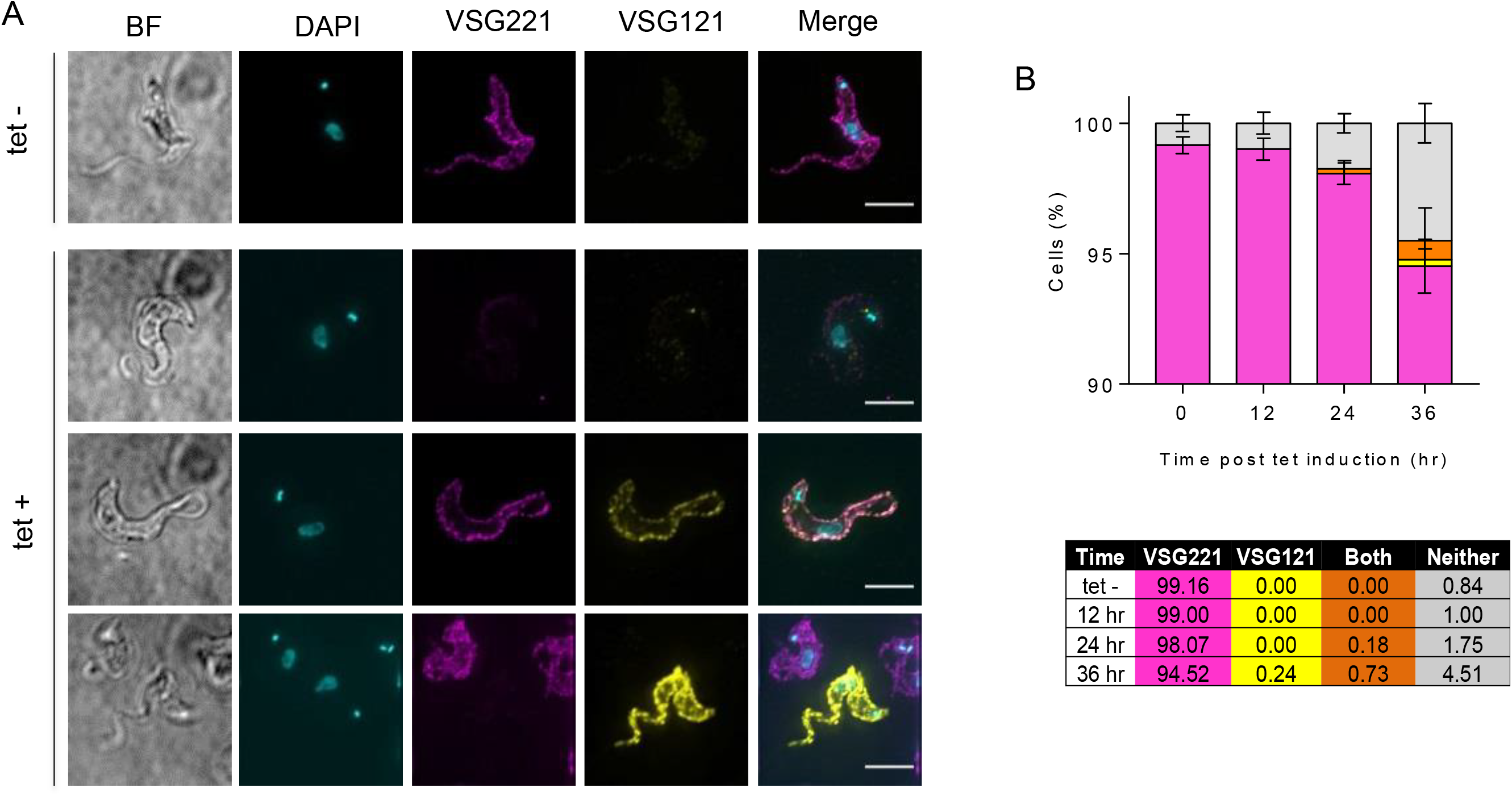
Depletion of TbRH2A induces switching of the VSG coat. A) Co-immunofluorescence imaging of VSG221 (pink) and VSG121 (yellow) surface expression. Example images are shown of cells after 24 hr of culture with (tet -) or without (tet +) RNAi induction (Scale bar, 5 μm). B) Graph of the percentage of uninduced (tet -, all time points) and RNAi-induced cells (after 12, 24 and 36 hr of culture with tet) expressing only VSG221 (pink) or only VSG121 (yellow) on their surface, as well as cell with both (orange) or neither (grey) of the two VSGs on their surface, as determined by coimmunofluorescence imaging with anti-VSG221 and VSG121 antiserum. >200 cells were analysed for each time point and three experimental replicates (error bars denote SEM). The table below shows the average percentage of the three replicates in each case.

## Discussion

RNase H enzymes that hydrolyse RNA within an RNA-DNA hybrid or remove ribonucleotides in DNA are ubiquitous in nature, with all organisms appearing to encode at least one RNase H, and most encoding two types of RNase H [26, 32]. Though such ubiquity suggests crucial roles, including in DNA replication, repair and transcription, loss of RNase H has so far only been described as being lethal during development in mammals [63] and *Drosophila melanogaster* [64]. In this study, we describe the roles played by RNase H2 in *T. brucei*, revealing the first example, to our knowledge, of an RNase H being essential in a single-celled eukaryote. The lethality we describe after TbRH2 loss contrasts with the non-essentiality of *T. brucei* RNase H1 [47, 52] and our data indicate that at least one explanation for this separation of function is that TbRH2 plays a specific and potentially novel role in processing R-loops at sites of multigenic RNA Pol II transcription initiation. Despite this distinction between the two *T. brucei* RNase H enzymes, we also reveal overlapping roles in targeting R-loops in RNA Pol I transcription units, contributing to the control and execution of immune evasion by antigenic variation [52]. Finally, gene expression analysis reveals changes in nucleotide and ribonucleotide synthesis and salvage pathways after TbRH2 loss, indicating that impairment of the parasite RNase H2 has cellular effects that parallel phenotypes described in humans with the autoimmune syndrome AGS [65], which can be caused by RNase H2 mutation.

RNAi-mediated depletion of TbRH2A led, within 24 hrs (approximately 3-4 cell cycles), to a growth arrest of BSF *T. brucei* cells that was accompanied by a pronounced impairment in the cell cycle, manifest initially as accumulation of 1N2K (G2/M phase) cells and then by the appearance of aberrant cells that failed to effectively divide their nucleus. Over the same period, TbRH2A RNAi resulted in increased expression of γH2A, which could be detected throughout the nucleus of most cells in the population, indicating more severe levels of nuclear genome damage than are seen after ablation of TbRH1 [52]. Unlike in yeast, where both RNase H genes can be deleted [33, 34], lack of continued proliferation of *T. brucei* cells after depletion of TbRH2A appears more comparable with truncated mammal embryogenesis seen in RNase H2B and RNase H2C mutants [63, 66]. Indeed, the nucleus-focused phenotypes after loss of RNase H2A in *T. brucei* are reminiscent of increased levels of γH2AX in cells from RNase H2 mutant mice [63, 66], and with fibroblasts taken from RNase H2B mutant mice showing slowed growth and accumulation in the G2/M phase of the cell cycle [66]. However, DNA content analysis revealed some differences when comparing RNase H2 mutant mouse cells and TbRH2A-depleted *T. brucei* parasites: flow cytometry and EdU incorporation analysis indicate that loss of TbRH2A does not prevent DNA synthesis, whereas RNase H2 mutant mice embryonic cells display reduced incorporation of EdU [63]. While this difference may only reflect variation between mouse and *T. brucei* cells in eliciting a cell cycle checkpoint in response to damage resulting from loss of RNase H2, perhaps due to changes in DNA damage signalling, it is also possible that genome replication is not the process primarily affected by TbRH2 loss in *T. brucei*. A less likely explanation, given the increased DNA content seen during flow cytometry in TbRH2A RNAi cells, is that *T. brucei*, unlike mice, enacts robust DNA repair synthesis, such break-induced replication, in response to lesions caused by loss of RNase H2.

Unlike RNase H1, RNase H2 has the capacity to excise ribonucleotides incorporated into the genome by initiating ribonucleotide excision repair [30, 32, 67], and lack of this activity has been proposed to underlie the embryonic lethality of mice RNase H2 mutants [63, 66]. However, yeast RNase H2 mutants also display increased levels of DNA-embedded ribonucleotides, leading to increased genome instability [33, 34], but survive. Equally, *D. melanogaster* RNase H1 mutants, which are likely to be unaltered in their capacity to remove ribonucleotides from the genome, display curtailed development during metamorphosis as a result of altered gene expression [64]. In this context, we suggest that the highly localised accumulation of γH2A and altered R-loop abundance we describe at sites of multigenic transcription initiation in the genome core after TbRH2A RNAi may provide an explanation for the importance of RNase H2 in *T. brucei*. We have shown previously that, amongst the widespread distribution of R-loops across the *T. brucei* genome, RNA-DNA hybrids display a clear association with the ∼200 mapped sites of RNA Pol II transcription initiation [47], with a strong correlation in localisation relative to epigenetic features found at such SSRs [42, 44, 68]. We now show that TbRH2A depletion results in pronounced γH2A accumulation only at these RNA pol II transcription initiation sites, not at termination sites or within the PTUs, and that these same loci are the only regions in the genome where DRIP-seq detects clear loss of R-loop abundance after RNAi. Strikingly, neither accumulation of damage nor loss of R-loops at the these sites is seen in TbRH1 null mutants [47]. The combined loss of R-loops and increased damage at these loci argue that these phenotypes are not the result of localised, increased levels of embedded ribonucleotides, but instead that RNase H2 has a specific role, not possessed by RNase H1, in processing R-loops associated with RNA Pol II transcription initiation. Why R-loop abundance might be reduced at transcription start sites after TbRH2 loss is unclear, but one explanation may be that the absence of RNase H2 allows inappropriate factors (perhaps even TbRH1) to attempt to remove the RNA-DNA hybrids, leading to damage [69]. Alternatively, failure to remove the R-loops due to loss of TbRH2 may stall RNA Pol II’s movement away from the start site, and the cross-linking we used to isolate R-loops might then occlude DRIP-seq signal. In these conditions, RNA pol II impediment of DNA replication [25], or activation of nucleotide excision repair due to stalling of RNA Pol [70], might lead to localised DNA lesions marked by γH2A.

Though there is clear evidence that R-loops are associated with genome instability, the processes that lead from an R-loop to DNA damage are less clearly understood [10, 71, 72]. Recently, Costantino and Koshland [73] mapped DNA damage in yeast cells impaired in R-loop removal, providing clear evidence for genome-wide correlation between sites of RNA-DNA hybrid formation and localisation of a damage marker, Rad52. Like we have described here, not all sites of R-loop formation in yeast lead to damage, indicating other processes much act to generate genome lesions. However, despite this similarity, significant differences are found in our data compared with the study by Costantino and Koshland [73]. First, damage accumulation in yeast requires mutation not only of both RNase Hs but also Senataxin or Topoisomerase I, whereas in *T. brucei* loss of only RNase H2 function leads to damage. Second, there was no evidence that the R-loop-associated damage in yeast shows the precise co-localisation with sites of transcription initiation that we have mapped in *T. brucei*. Thus, though it is conceivable that the damage we see, which was detected via γH2A, might also be sites of single-stranded DNA bound by Rad52 [73], distinct routes of damage formation are likely. Intriguingly, mounting evidence has linked the generation of DNA breaks and the action of DNA repair factors in transcriptional activation [74–76], including regulating elongation of RNA Pol at mammalian protein-coding and non-coding RNA genes [77, 78], and altering chromatin to activate gene expression during *Caenorhabditis elegans* embryogenesis [79]. Thus, our data suggest it is possible that DNA breaks are also a feature of transcription initiation in *T. brucei*, with their extent or persistence increased by loss of RNase H2. Mapping RNA Pol II by ChIP has revealed accumulation at transcription start sites, consistent with pausing [44], which correlates with R-loops at these sites [47]. It is important to note, however, that there is no evidence for *T. brucei* RNA Pol II transcription being controlled at the point of initiation [44], so whether the precise association we observe between RNase H2-associated DNA damage and R-loop levels at the start of multigenic transcription in *T. brucei* might have parallels with single gene, regulated transcription in other eukaryotes is currently unknown. Nonetheless, R-loops are readily detected at CpG island promoters in the human and mouse genomes [80, 81]. Moreover, near genome-wide multigenic RNA Pol II transcription is common to all kinetoplastids [39], meaning it is likely that RNase H2 plays related roles in transcription initiation throughout this grouping of microbes and may be fertile ground for new therapies against diseases caused by the parasites.

The RNA-seq we describe here evaluates differential expression of gene-specific transcripts before and after TbRH2A RNAi, which reveals two aspects of TbRH2 function. First, as discussed more fully below, the genes most rapidly and most strongly affected by loss of TbRH2 are transcribed by RNA Pol I, not RNA Pol II. Second, amongst the cohort of differentially expressed RNA Pol II transcripts, we reveal parallels with emerging understanding of human disease caused by loss of RNase H2. AGS has emerged as a valuable model for complex autoimmune disorders because the disease can be caused by a single gene mutation [65], including in all three subunits of RNase H2 as well as several other genes that encode enzymes with nucleic acid or nucleotide targeting activities [35]. Mouse models of AGS carrying RNase H2 mutations have shown that activation of the cGAS-STING immune sensing pathway elicits a type I interferon response associated with AGS pathological symptoms [36, 37]. Cyclic GMP-AMP synthase (cGAS) normally detects cytosolic DNA via the cyclic GMP-AMP receptor STING, mediating one arm of the innate immune response against pathogens [82]. Since the absence of fully functional RNase H2 leads to AGS, it is considered likely that increased abundance of DNA with embedded ribonucleotides, or RNA-DNA hybrids could aberrantly activate the cGAS-STING pathway [36, 37]. Over-expression of RNase H1 in RNase H2B null mutant mice is unable to reduce DNA damage levels or interferon simulated gene expression to WT levels, implicating DNA with embedded ribonucleotides as the main nucleic acid activating the immune response associated with AGS [37, 83]. However, it remains possible that aberrant breakdown and release of some forms of RNA-DNA hybrid, which RNase H1 may be unable to efficiently resolve, could also induce the cGAS-STING pathway. In *T. brucei* we have identified a specific activity of RNase H2 at RNA Pol II transcription initiation sites, a role which *T. brucei* RNase H1 appears not to perform [47]. RNA-seq analysis of TbRH2A depleted parasites revealed downregulation of nucleotide and ribonucleotide synthesis and salvage pathways, which was not seen when RNA-seq compared gene expression in *TbRH1* null and WT parasites [47], suggesting increased levels of dNTP and rNTP pools in the parasites. A key question in the development of AGS is the source and form of the host cytoplasmic nucleic acid that causes autoimmunity. The RNA-seq profiles we describe seem to indicate that loss of *T. brucei* RNase H2, but not RNase H1, releases nucleic acids, leading to altered gene expression, perhaps as part of wider metabolic regulation. Such an effect may, like in AGS, be due to RER, but may also relate to the locus-specific damage and aberrant R-loop processing during transcription we describe after TbRH2A loss, which may also occur during AGS but has not yet been examined.

Despite the pronounced, localised effect of TbRH2A RNAi at sites of RNA Pol II transcription initiation, RNA-seq revealed that a stronger and earlier effect was exerted on protein-coding genes expressed by RNA Pol I. Most notably, TbRH2A loss resulted in increased RNA levels of previously silent VSGs, leading to changes in the surface coat. As these changes in VSG expression were concomitant with increased accumulation of R-loops and DNA damage in the telomeric VSG BES, and reflect similar findings described in *TbRH1* null mutants [52], the data provide further evidence that R-loops acted upon by RNase H are an important driver of antigenic variation through VSG switching. Similarities and differences in the VSG-associated phenotypes seen after TbRH2A RNAi and ablation of TbRH1 suggest overlap in the function of the two enzymes in this reaction, but also some divergence. In both TbRH2A-depleted and *TbRH1* null cells, R-loop levels appeared to increase to similar levels in both the active and silent BES. Though these data suggest that loss of either RNase H impairs resolution of BES R-loops and leads to VSG switching, how the RNA-DNA hybrids accumulate in the silent ES, which are not transcribed to the same extent as the main active ES, remains unclear. Despite this similarity, levels of BES damage measured by γH2A ChIP differed: whereas loss of TbRH1 results mainly in increased damage towards the telomere of the active BES, TbRH2A depletion caused greater amount of yH2A accumulation throughout both the active and silent BES. This difference may reflect a greater impact of TbRH2A depletion on transcription, although greater damage induction due to persistence of ribonucleotides in the BES cannot be ruled out; indeed, increased incorporation of uracil into DNA, due to loss of uracil-DNA glycosylase, has been shown to lead to DNA lesions and VSG switching [84]. Nonetheless, two pieces of evidence suggest that loss of TbRH2A affects BES transcription more that ablation of TbRH1. First, whereas RNA-seq only revealed increased expression of VSGs in *TbRH1* mutants, increased levels of RNA from promoter-proximal ESAGs, as well as from ES VSGs, was detected by the RNA-seq described here. Second, in TbRH2A depleted parasites, significantly higher numbers of cells were found that expressed two VSGs simultaneously on their surface compared with the same IF analysis of *TbRH1* null mutant cells. Hence, loss of RNase H2 compromises more strongly the strict monoallelic expression of VSG BES normally employed by *T. brucei* [85]. Whether this is because the TbRH2 complex recruits factors involved in monoallelic control, while TbRH1 does not, is worthy of investigation; for instance, it is known that RNase H2B in other eukaryotes interacts with PCNA [86]. We have argued that DNA damage induced in the BES by R-loops, and accentuated by loss of TbRH1, is a plausible route for the initiation of recombination of silent VSGs [52]. This argument is supported here by the finding that TbRH2A RNAi also led to increased BES damage and expression of silent VSGs from throughout the repertoire, including intact subtelomeric genes, minichromosome genes and pseudogenes, which are unlikely to be transcribed without movement to the BES. Nonetheless, how, when and where BES R-loops are converted into DNA damage, and the nature of the lesions that arise, remain open questions.

## Methods

### *T. brucei* cell line generation and culture

C-terminal endogenous epitope tagging was carried out by cloning 604 bp of the 3’ *TbRH2A* sequence, PCR-amplified using primers CGACGAAGCTTGAACACGCTTAGCCATCAAA*C* and GACGTCTAGAAGGGACTTCCCGCGACAAA, into a version of the pNAT plasmid containing 6 copies of the HA sequence [87]. The construct was linearized by digestion with ApaI, before stable transfection into BSF *T. b. brucei* strain Lister 427 MITat1.2, where the construct was integrated immediately downstream of the *TbRH2A* coding sequence. Inducible RNAi targeting TbRH2A was accomplished using a genetically modified strain derived from Lister 427 MITat1.2, named 2T1 [88]. Two inverted copies of 432 bp of the *TbRH2A* ORF, PCR-amplified using primers GGGGACAAGTTTGTACAAAAAAGCAGGCTCCGCAGCTATGACAGGTGTA and GGGGACCACTTTGTACAAGAAAGCTGGGTCTCGAAGACGATAGGGATG, were cloned into the pGL2084 vector [57] using Gateway BP Clonase. The construct was then linearised and transfected into the parental 2T1. Incubation of the resulting TbRH2A^RNAi^ cell line with 1 μg.mL^−1^ tetracycline induced transcription of the *TbRH2A*-specific hairpin RNA molecules, triggering RNAi. Lister 427 and 2T1 cells, as well as transformants derived from each, were maintained in HMI-9 (HMI-9 (Gibco), 20 % v/v FCS (Gibco), Pen/Strep (Sigma) (penicillin 20 U/ml, streptomycin 20 μg/ml)), and HMI-11 (HMI-9 (Gibco), 10 % v/v FCS (Gibco, tet free), Pen/Strep (Sigma) (penicillin 20 U/ml, streptomycin 20 μg/ml)) media, respectively. During EdU uptake assays, cells were instead incubated in thymidine-free HMI-11 media (Iscove’s Modified Dulbecco’s Medium (IMDM) (Gibco), 10% (v/v) FBS (Gibco, tet free), 1% (v/v) Pen/Strep (Gibco), 4% (v/v) “HMI-9 mix” (0.05 mM bathocuproine disulphonic acid, 1 mM sodium pyruvate and 1.5 mM L-cysteine) (Sigma Aldrich), 1 mM hypoxanthine (Sigma Aldrich) and 0.0014% 2-mercaptoethanol (Sigma Aldrich)). In all cases, 2T1 cells lines were grown in the presence of 0.5 μg.mL^−1^ puromycin (InvitroGen) and 2.5 μg.mL^−1^ phleomycin (InvitroGen), and TbRH2A^RNAi^ parasites were cultured with 2.5 μg.mL^−1^ phleomycin (Invitrogen) and 5 μg.mL^−1^ hygromycin B (Calbiochem) to maintain the RNAi construct.

### Fluorescent-activated cell sorting (FACS)

∼3 x10^6^ cells were collected per sample and fixed with 1% formaldehyde (FA) for 10 min at room temperature before being permeabilized with 0.01 % Triton X-100 for 30 min on ice. Cells were incubated with 100 μg.mL^−1^ RNase A and 15 μg.mL^−1^ propidium iodide (PI) for 30 min before PI was detected using the BD FACSCalibur™ (BD Biosciences). Data was analysed using FlowJo_V10™ software (FlowJo, LLC).

### Immunofluorescence imaging

All immunofluorescence assays were performed as previously described [52]. For detection of epitope-tagged TbRH2A and γH2A, cells were fixed with 4% FA for 4 min before quenching with 100 nM glycine and permeabilization with 0.2% Triton X-100 for 10 min. 3% bovine serum albumin was using to block samples before staining with primary (α-myc Alexa Fluor 488 conjugated (Millipore), 1:500; α-γH2A, 1:1000) then secondary antibodies (Alexa Fluor 488 goat α-rabbit (Molecular Probes), 1:1000) and mounting in Fluoromount G with DAPI (Cambridge Bioscience, Southern Biotech). For VSG immunofluorescence cell were fixed in 1% FA and blocked in 50% foetal calf serum, before staining with primary (α-VSG221, 1:10,000; α-VSG121, 1:10,000: gift from D. Horn) and secondary (Alexa Fluor 594 goat α-rabbit (Molecular Probes) 1:1000; Alexa Fluor 488 goat anti-rat (Molecular Probes), 1:1000) antibodies and DAPI mounting as above. Fluorescent imaging was performed with an Axioscope 2 fluorescence microscope (Zeiss) and a Zeiss PlanApochromat 63x/1.40 oil objective. High-resolution images were taken using an Olympus IX71 DeltaVision Core System microscope (Applied Precision) and SoftWoRx Suite v2.0 (Applied Precision) software. Either an Olympus PlanApo 60x/1.42 or an UplanSApo 100x/1.40 oil objective was used. Z-stacks of 5-6 μm thickness were acquired in 25 sections, then deconvolved with the ‘conservative’ method and high noise filtering. Fiji software was then used to generate maximum projection images. Super-resolution structured illumination microscopy (SR-SIM) was performed with an ELYRA PS.1 Microscope (Zeiss), using a Plan-Apochromat 63x/1.40 Oil DIC objective and 405, 488 and 594 nm lasers. Z-stack sections were ∼0.15 μm in thickness and totalled ∼10 μm. Image reconstruction was performed with ZEN Black software (Ziess) and 3D rendering with Imaris software (Bitplane) to produced 3D models.

### Western blot

Whole cell protein extracts were harvested from ∼ 2.5 x 10^6^ cells per sample by boiling in 10 μl loading buffer (1X NuPAGE^®^ LDS Sample Buffer (Life Technologies), 0.1% β-mecaptoletanol, 1X PBS) for 10 min. Proteins were separated using NuPAGE Novex^®^ 10-12% Bis-Tris protein gels (Life Technologies) and transferred to PVDF membranes. Proteins were detected with anti-γH2A (1:1000) and mouse anti-Ef1α clone CBP-KK1 (1:25,000; Millipore) primary antibodies, and goat anti-mouse/rabbit IgG (H+L) horseradish peroxidase conjugates (1:5000; Life Technologies).

### EdU incorporation assays

Cells were incubated with 150 μM 5-ethynyl-2′-deoxyuridine (EdU) for 4 hr prior to fixation at each time point with 1% FA at room temperature for 10 min, then permeabilised in 0.5% Triton X-100 for 20 min. EdU was detected by incubation for 1 hr with the follow Click-It reaction mix: 21.5 μl 1X Reaction Buffer, 1 μl CuSO4, 0.25 μl Alexa Fluor 555 Azide and 2.5 μl 1X Additive Buffer. For dual staining for γH2A, cells were washed before incubating with γH2A antisera (1:1,000) then anti-rabbit Alexa Fluor 594 (1:1,000), both in 3% BSA, and mounted in Fluoromount G with DAPI.

### DRIP/ChIP analysis

Immunoprecipitation of both RNA-DNA hybrids (DRIP) and γH2A was performed using the ChIP-IT Enzymatic Express kit (Active Motif) with formaldehyde fixed chromatin samples, using the hybrid-targeting S9.6 (4.5 ng, Kerafast) and α-γH2A (10 μl, homemade) antibodies respectively. Both were carried out as described previously [47]. DRIP-ChIP-qPCR analysis was carried out as described previously [52] with on-bead *E. coli* RNase HI treatment of DRIP samples.

DNA libraries were prepared using the TruSeq ChIP Library Preparation Kit (Illumina). Fragments of 300 bp, including adaptors, were selected with Agencourt AMPure XP (Beckman Coulter) and sequenced using the Illumina NextSeq 500 platform. Analysis of DRIP-seq and γH2A ChIP-seq data was analysed as previously published [47]. To allow downstream analysis to focus on the γH2A signal specific to the TbRH2A depletion induced DNA damage, the ratio of read enrichment was calculated for tet-induced samples relative to un-induced sample coverage (having first normalised to the respective input sample) for both 24 and 36 hr timepoint data sets. All normalisation and metaplot analysis was performed with the deepTools software suite [89].

### RNA-seq analysis

Total RNA was extracted, in duplicate, from 1 x 10^7^ cells using the RNeasy Mini Kit (Qiagen) and the TruSeq Stranded mRNA kit (Illumina) was used to prepare poly(A) selected libraries of ∼300 bp fragments. Sequencing was performed with the NextSeq 500 platform giving paired-end reads of 75 bp. Mapping was performed with HiSat2 v2.0.5 using default parameters with the exception of not permitting splice alignments (--no-splice-alignment), to either a “hybrid” reference genome, consisting of the 11 Mb chromosome assemblies of the *T. brucei* TREU927 v5.1 genome, 14 BES contigs and 5 mVSG ES contigs [61], or a collection of 2470 VSG coding regions of the *T. brucei* Lister427 strain [62]. Reads with MAPQ score <1 were removed before counting with HTseq-count software using default parameters. Normalisation and differential expression was carried out with DESeq2 v1.18.1 [90], considering data from 24 and 36 hr time points separately. GO term analysis was performed using Cytoscape v3.6.1 [91] plugin BiNGO [92] with hypergeometric statistical testing of significance and multiple testing correction with the Benjamini and Hochberg False Discovery Rate (FDR) correction. FDR adjusted P values < 0.01 were deemed significantly enriched terms.

## Acknowledgements

We thank all lab members for invaluable discussions, and Sebastian Hutchinson for providing sequences and comparing R-loop data. This work was supported by the BBSRC [BB/K006495/1,BB/M028909/1, BB/N016165/1 and a DTP studentship to E. Briggs]. The Wellcome Centre for Molecular Parasitology is supported by core funding from the Wellcome Trust [104111].

## Author contributions

EB, KC and RM designed the experimental strategy. EB, LL and CL conducted the molecular and imaging experiments. EB, KC, GH and RM analysed and interpreted next generation mapping data. EB, KC, LL, GH and RM wrote the paper. All authors provided intellectual input and approved the final manuscript.

## Data Access

Sequences used in the mapping have been deposited in the European Nucleotide Archive (accession number PRJEB22057).

## Supplementary figure legends

**Table S1.**
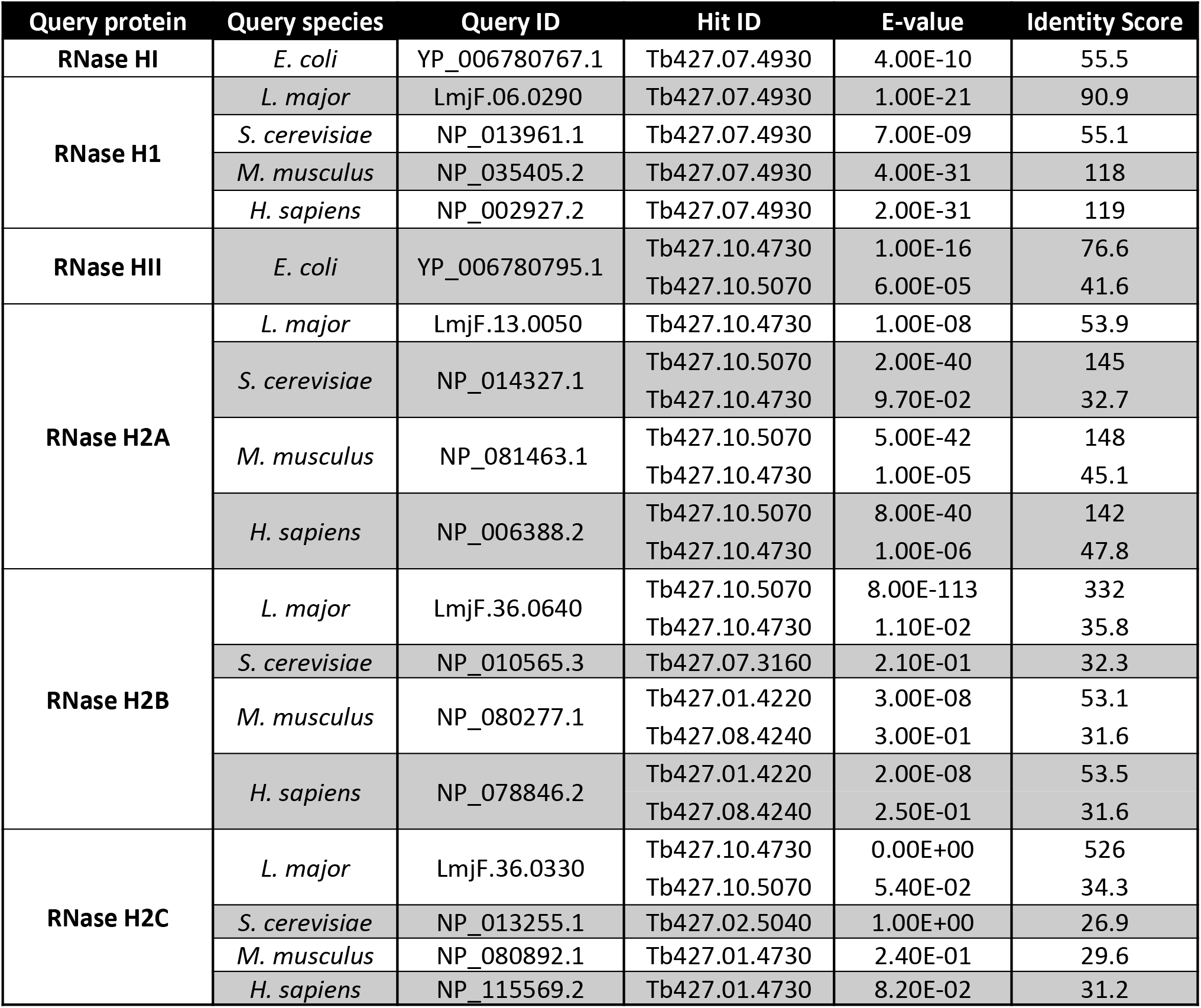
Identification of genes encoding class 1 and class 2 RNase H enzymes in the genome of *Trypanosoma brucei*.

**Figure S1.**
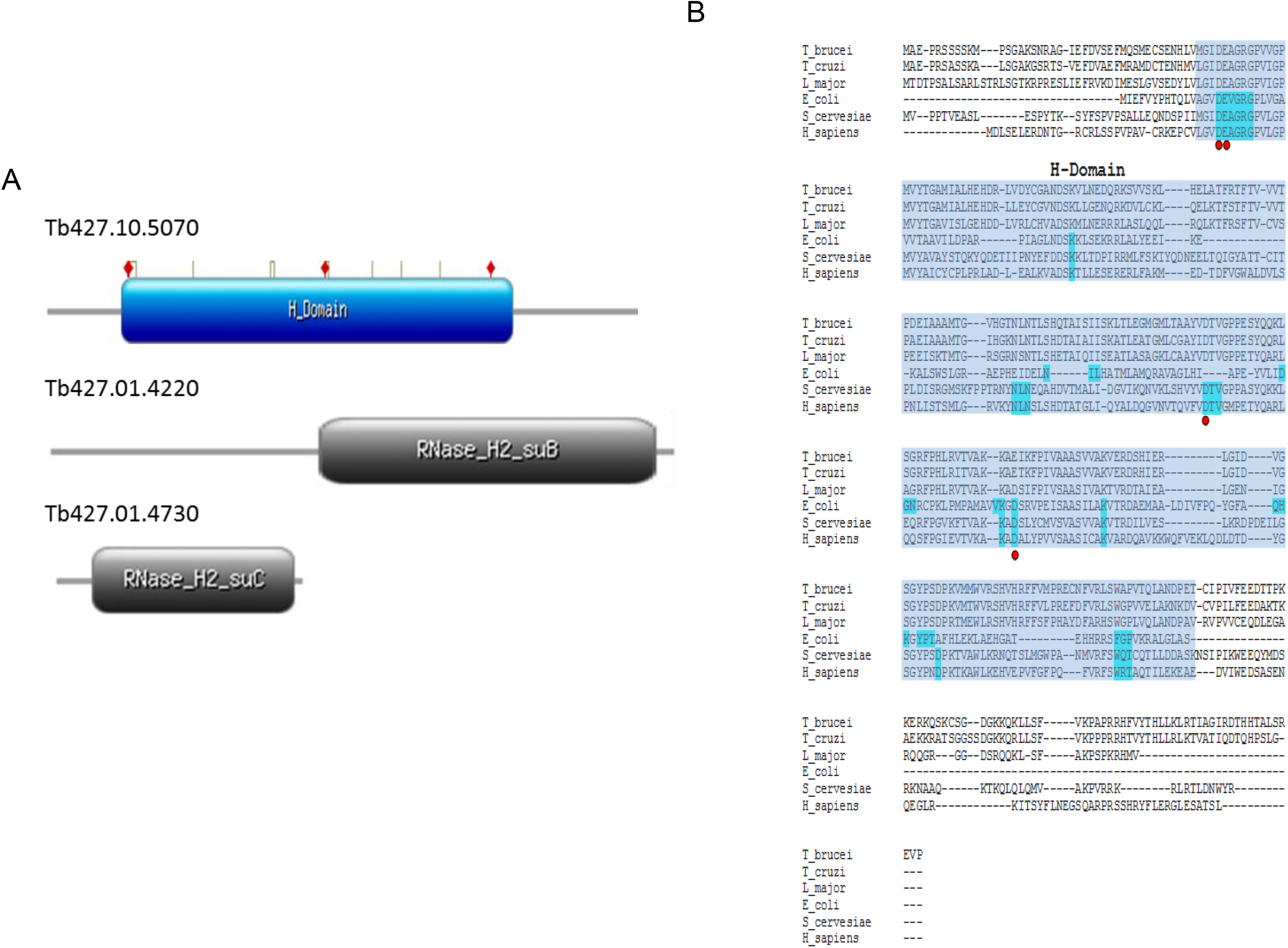
Protein sequence analysis of putative *T. brucei* type 2 RNase H proteins. A) Structural models of putative RNase H2 subunits A (Tb427.10.5070), B (Tb427.01.4220) and C (Tb427.01.4730), as predicted by InterPro analysis. Conserved domains are shown, and active site and catalytic residues are highlighted as white boxes and red diamonds above predicted domains, respectively. B) Amino acid alignments of putative RNase H2A catalytic proteins from *T. brucei, T. cruzi, L. major, E. coli* and *T. maritina*. Predicted sequences of the RNase H (H-)domain is highlighted in blue, active site residues in cyan, and catalytic residues are indicated by red dots.

**Figure S2.**
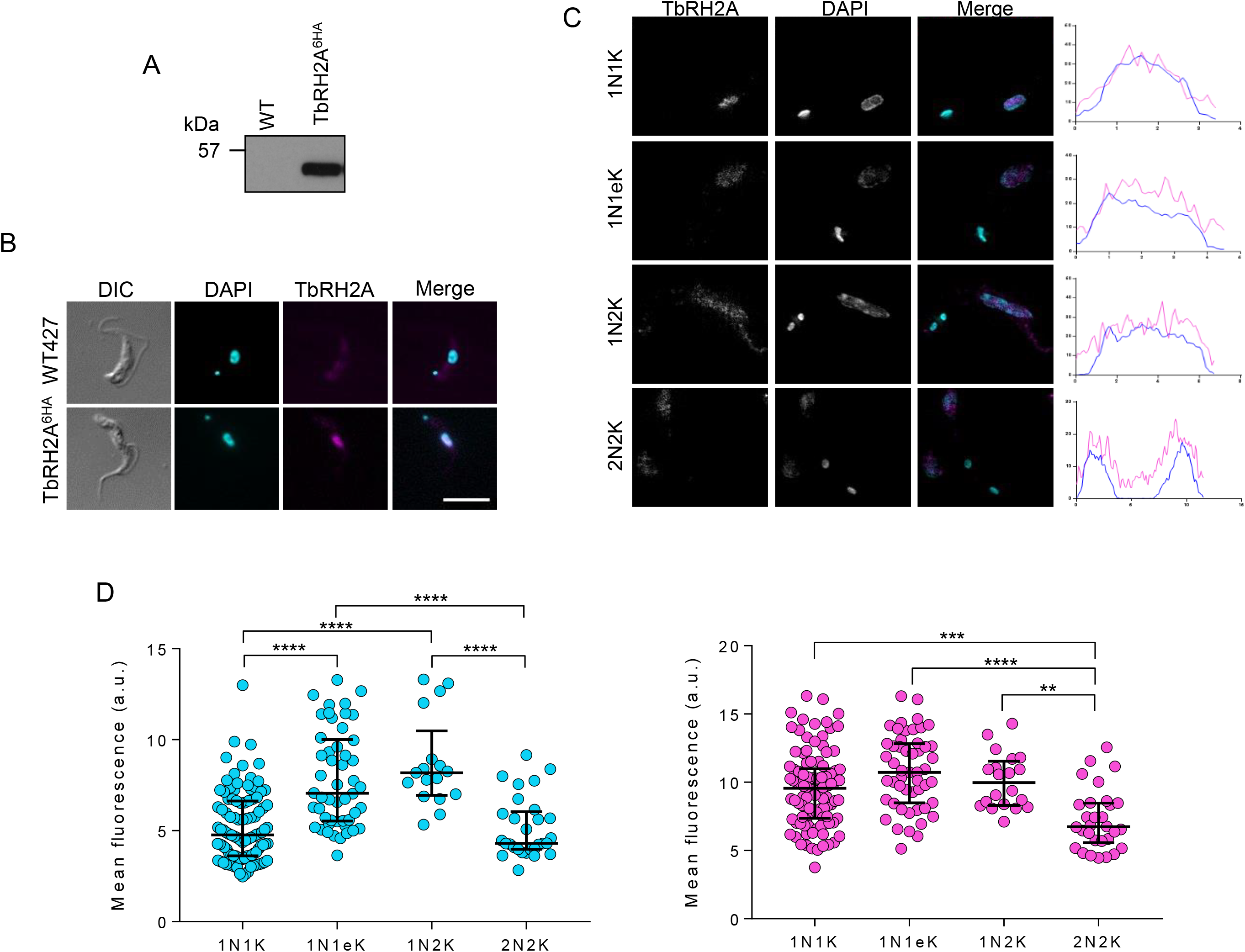
TbRH2A is a nuclear protein in bloodstream form *T. brucei* parasites. A) Western blot detection, using anti-HA antiserum, of C-terminally 6HA-tagged TbRH2A protein expressed from its endogenous locus in clonal bloodstream form *T. brucei*. A sample from untagged wild-type (WT) parasites is shown for comparison along with the expected protein size (kDa). B) Representative immunofluorescent images of *T. brucei* parasites expressing 6HA-TbRH2A; WT cells are shown for comparison. Anti-HA signal is shown in magenta and DAPI staining in cyan. The cell outline is shown by differential interference contrast microscopy (DIC). Scale bar, 5μm. C) Super-resolution structured illumination imaging of TbRH2A and nuclear DNA (magenta and cyan respectively in the merged, uncoloured images), detected with anti-HA antiserum and DAPI respectively. A respective cell of each cell cycle stage is shown along with graphs plotting length across the nucleus (x-axis, pixels) versus mean pixel intensity at each point (y, arbitrary units) for DAPI (cyan) and TbRH2A (magenta). Scale bars, 5μm. D) Graphs depicting mean fluorescence intensity (arbitrary units, a.u.) of DAPI (left, cyan) and anti-HA-staining (right, magenta) to detect TbRH2A expression. Cells are separated by discernible cell cycle stages (determined by number of nuclear (N) and kDNA (K) signals visualised by DAPI staining; 1N1K, 1N1elongatedK (1N1eK), 1N2K and 2N2K). Each data point denotes the mean fluorescence of an individual cell and the median values (horizontal lines) and interquartile range (error bars) are shown. Significance was determined by the Kruskal-Wallis non-parametric test: (**) p-value <0.01, (***) p-value <0.001, (****) p-value < 0.0001.

**Figure S3.**
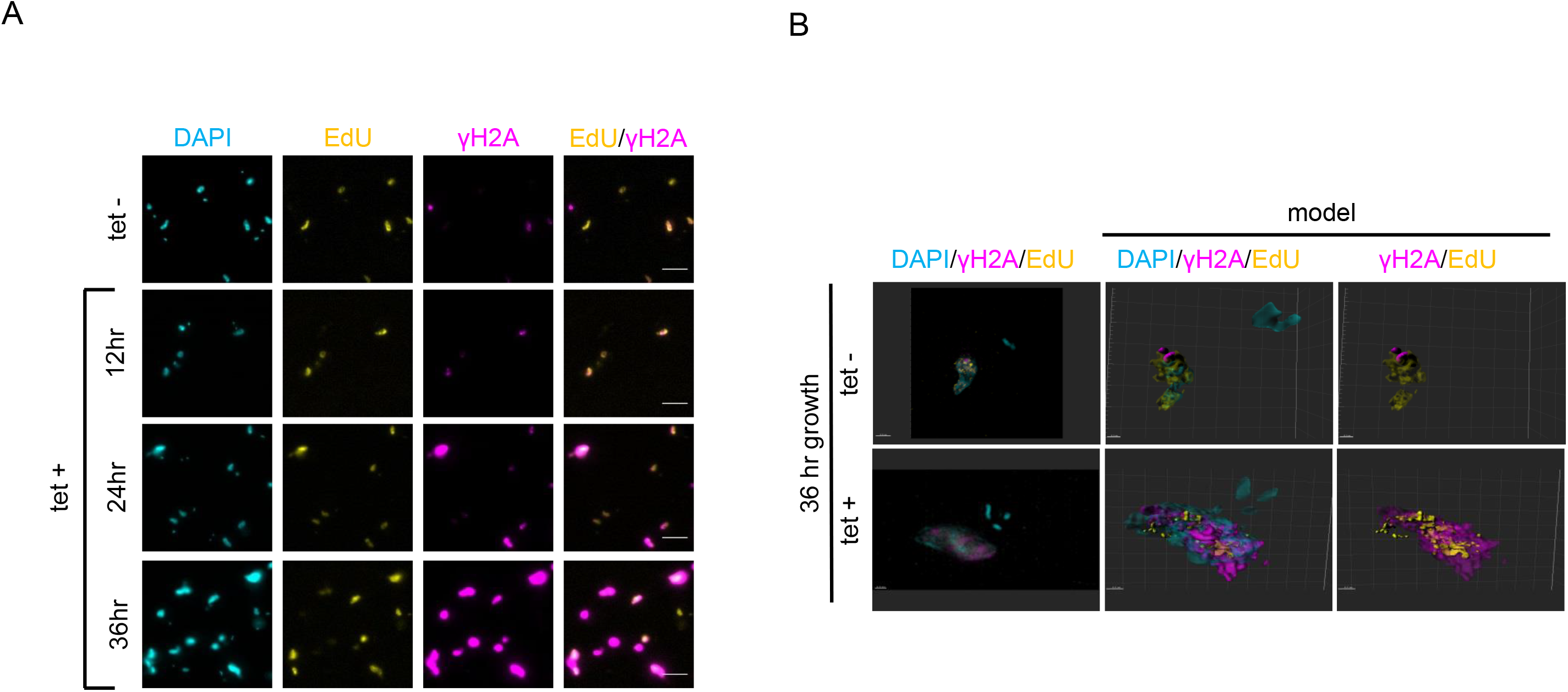
DNA synthesis occurs in the presence of DNA damage upon depletion of TbRH2A in *T. brucei*. A) Immunofluorescence detection of nucleic acid (DAPI staining; cyan), thymidine analogue EdU incorporation (via Click-IT chemisty; yellow) and γH2A (anti-γH2A antiserum; magenta) in TbRH2A RNAi parasites grown in either the absence (tet -) or presence of tet-induction (tet +) for 12, 24 or 36 hr. EdU and γH2A signal colocalization within several cells is also shown as merged images. Scale bars, 5μm. B) SR-SIM images of DAPI, γH2A and EdU staining are shown, along with 3D reconstructions (model), of TbRH2A RNAi cells grown in the absence (tet -) or presence (tet +) of tet-induction. Scale bars, 1 μm.

**Figure S4.**
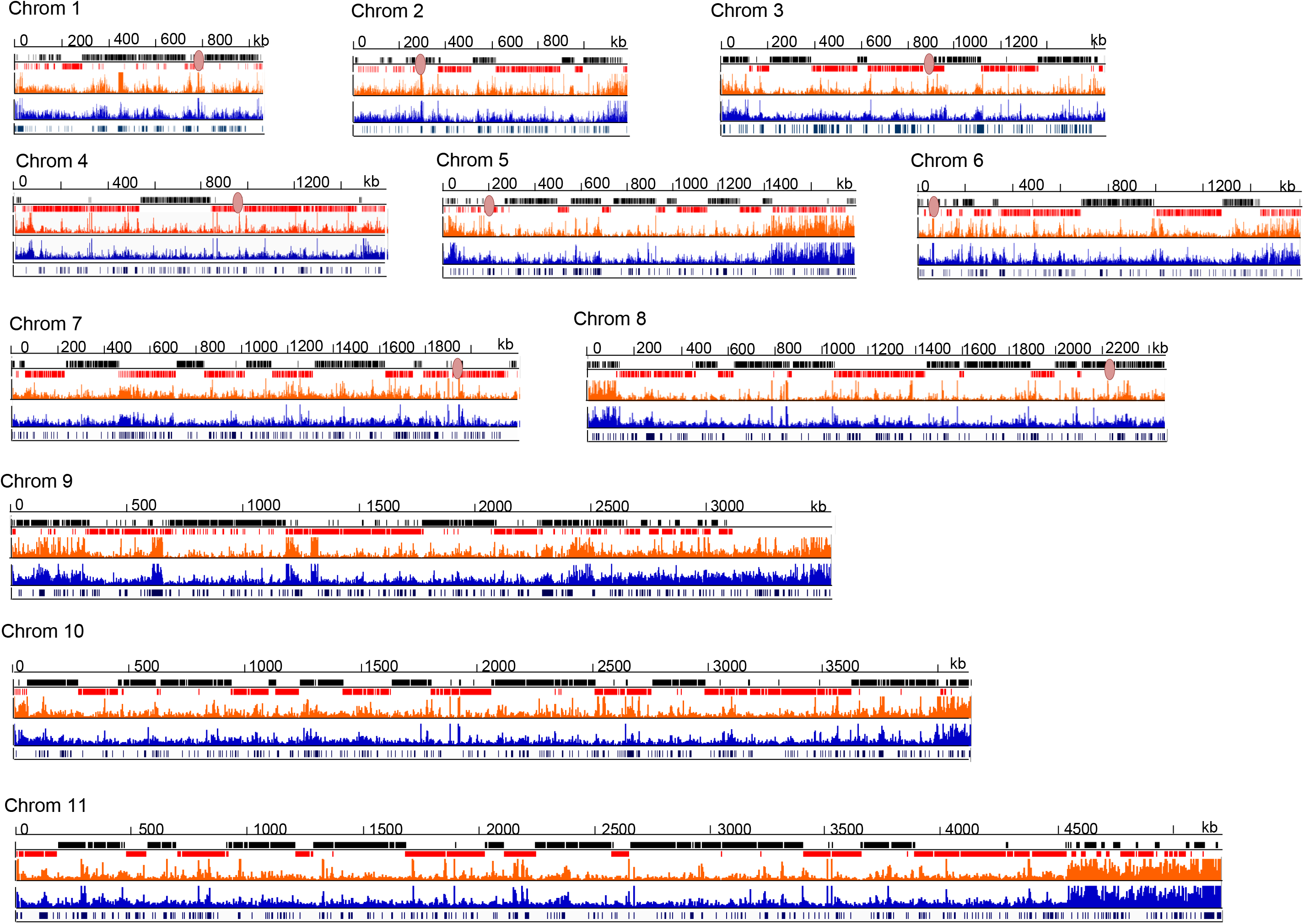
DRIP-seq signal across the 11 Mb-sized chromosomes of *T. brucei* before and after TbRH2A depletion. DRIP-seq signal for TbRH2A RNAi parasites grown without (blue) and with (orange) tet-induction for 24 hr is plotted as fold-change of IP sample coverage relative to input sample coverage (scale: 1-4 fold change). Upper track shows genes encoded in the sense (black) and antisense (red) directions. Known centromeres are shown as peach ovals. Lowest track shows predicted tandem repeat sequences.

**Figure S5.**
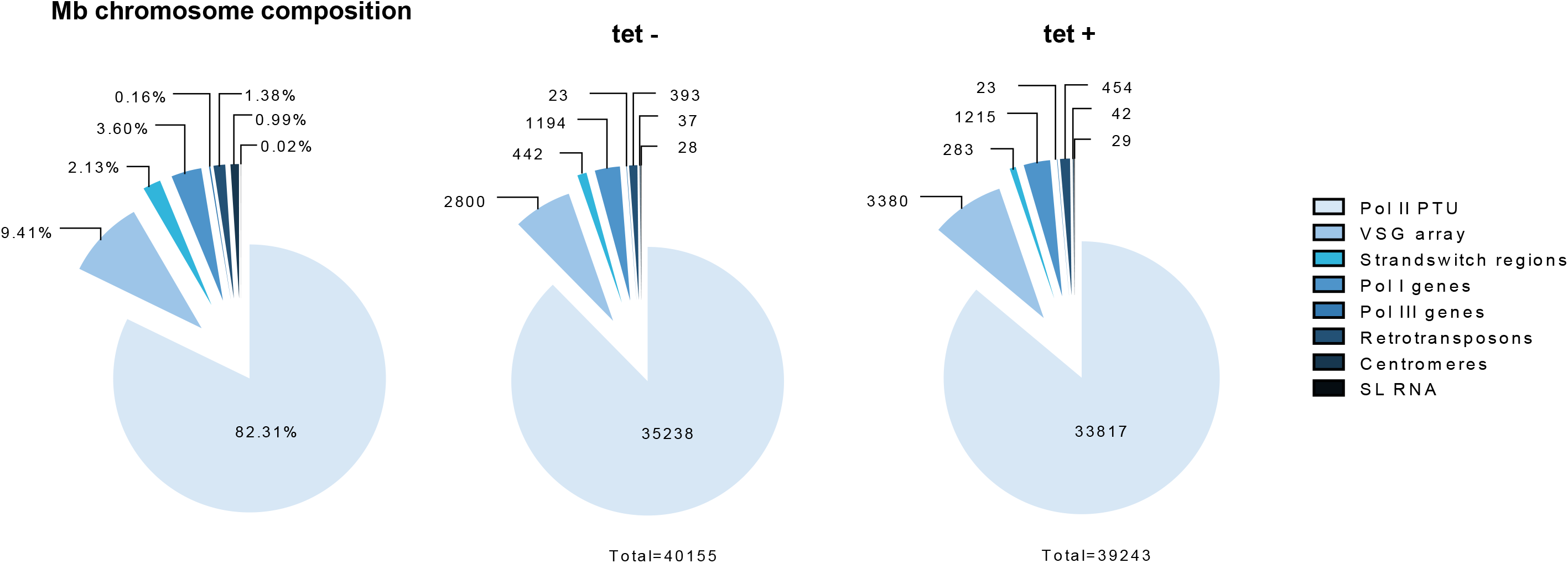
Distribution of DRIP enriched regions across the 11 Mb-sized chromosomes. The distribution of DRIP-seq enriched regions (> 1.2 fold change DRIP signal relative to input signal) in TbRH2A RNAi tet-induced (tet +) and uninduced (tet -) parasites. The composition of the Mb-sized chromosomes is shown to the left for comparison.

**Figure S6.**
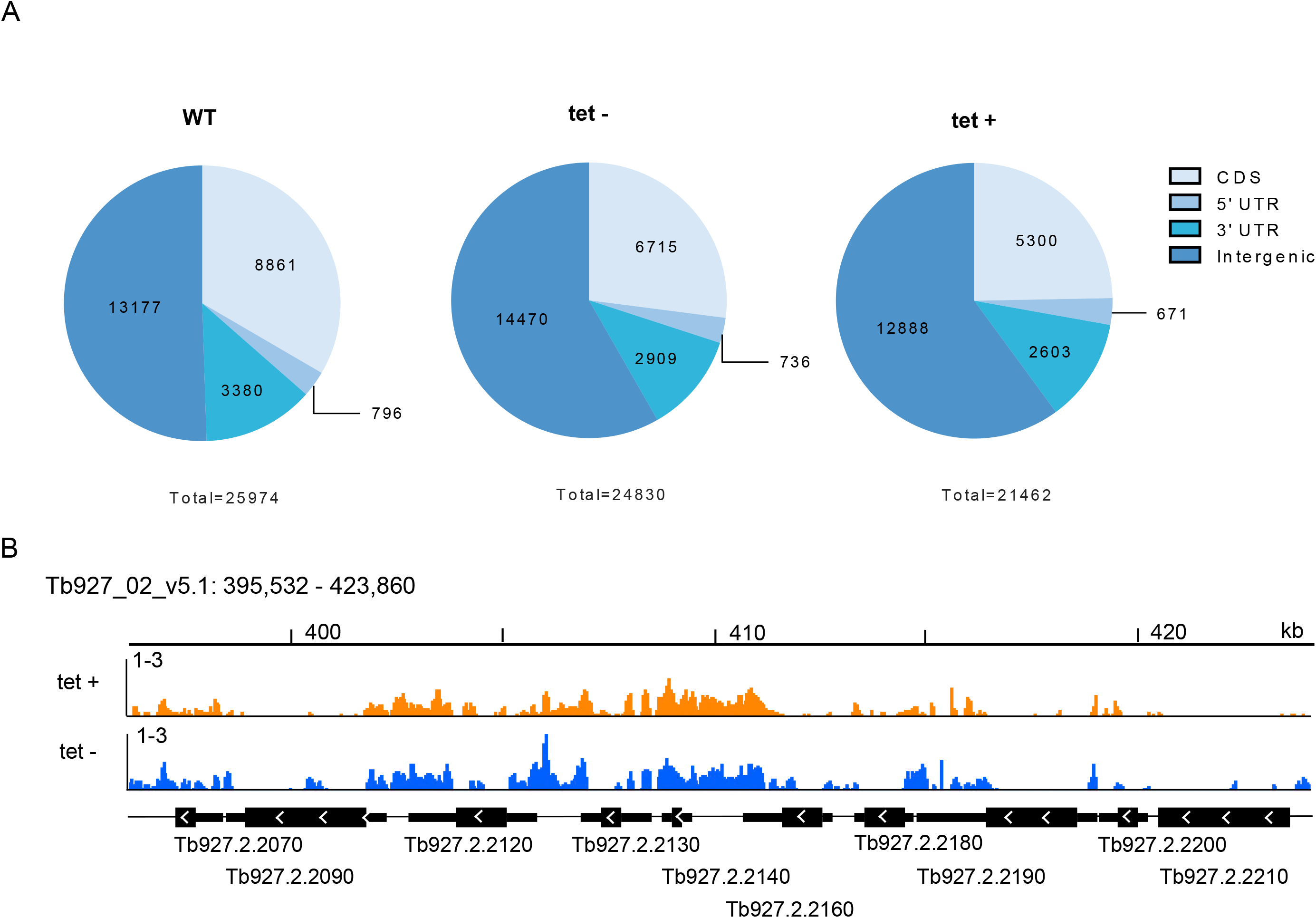
Distribution of DRIP enriched regions across the RNA Pol II transcribed polycistronic units. A) The distribution of DRIP enriched regions between CDS, 5’ UTR, 3’ UTR and intergenic sequences within the RNA Pol II transcribed PTUs is shown for WT, TbRH2A RNAi uninduced (tet -) and induced (tet +) cells. Total numbers of DRIP enriched regions defined are displayed below plots. B) DRIP-seq signal coverage (normalised to input sample coverage; scale 1-3 fold-change) across an example regions of a PTU within Mb chromosome 2, for TbRH2A RNAi uninduced (tet -) and induced (tet +) samples. CDS regions are indicated by thick black lines below tracks, UTR sequences are indicated by thin black lines, and white arrows indicate transcription direction.

**Figure S7.**
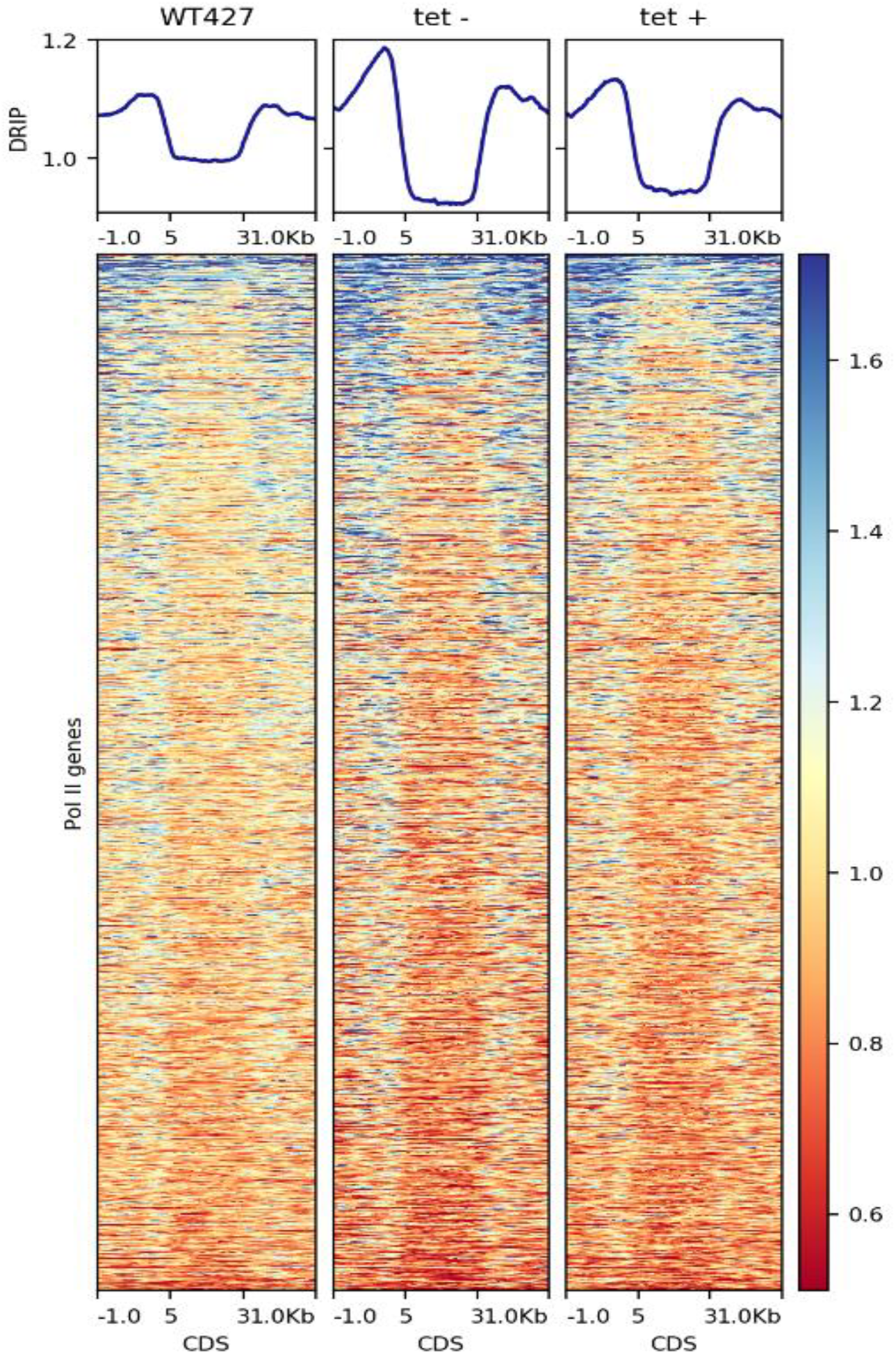
DRIP-seq signal is enriched in regions flanking CDS of RNA Pol II transcribed genes. Heatmaps of DRIP-seq signal (normalised to input signal) across each RNA Pol II transcribed protein-coding CDS, plus 1 kb of upstream and downstream flanking region, are shown for WT, TbRH2A RNAi uninduced (tet -) and induced (tet +) cells. Profiles of the average DRIP-seq of all CDS is shown above each heatmap.

**Figure S8.**
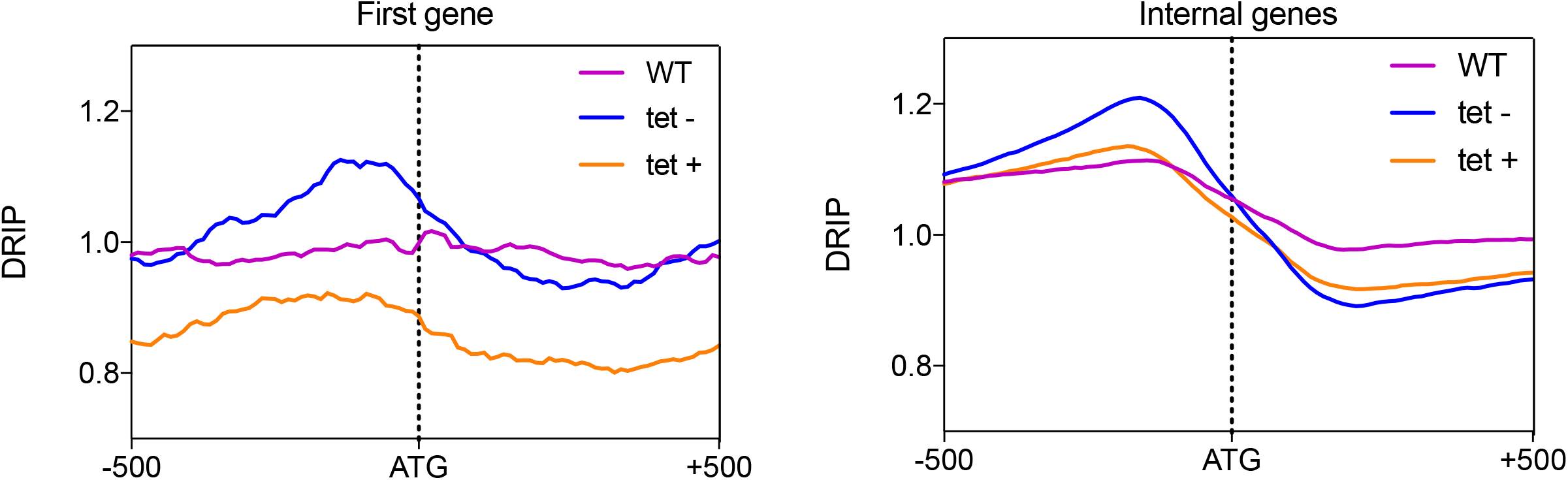
DRIP-seq signal mapping across the translational start sites of RNA Pol II transcribed genes. Average DRIP-seq signal coverage (normalised to input coverage; x axes) is plotted across 1 kb surrounding the ATG start sites (y axes) of the first gene in each RNA Pol II transcribed PTU (left), and all other genes contained with the PTUs (right). Average signal is shown for WT (pink), TbRH2A RNAi uninduced (blue; tet -) and induced (orange; tet +) cells. Shaded areas show SEM.

**Figure S9.**
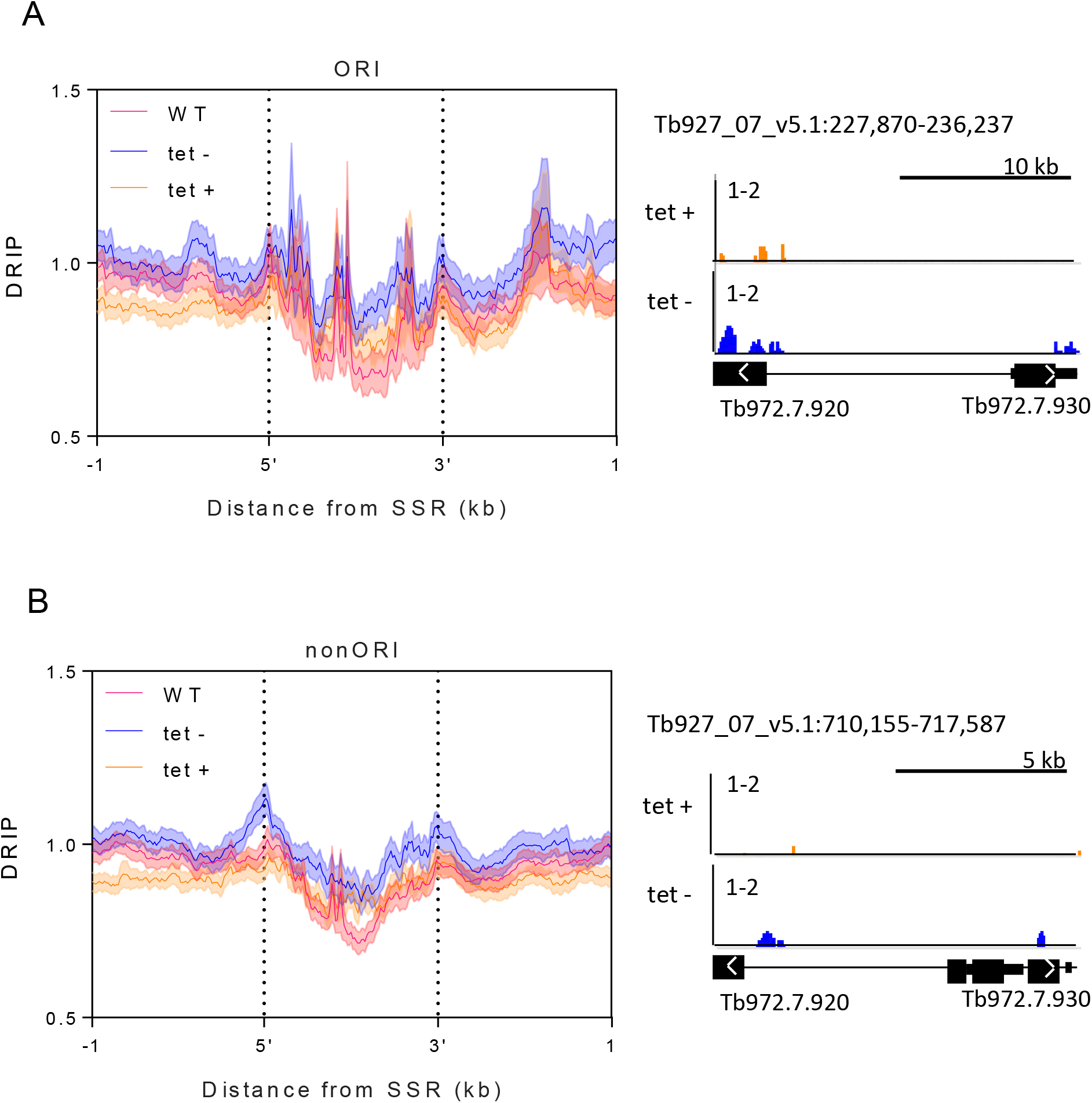
Levels of DRIP-seq signal are not altered in DNA replication origins SSRs compared with non-origin SSRs. A) The average DRIP-seq signal coverage (normalised to input sample coverage) is plot across SSRs, plus 1 kb of upstream and downstream flanking regions, which are known origins of replication (ORI) for WT (pink), TbRH2A uninduced (blue; tet -) and induced (orange, tet +) cells. The 5’ and 3’ boundaries of each SSR were defined as the end and start of flanking transcripts. Shaded areas represent SEM. An example of DRIP-seq signal coverage is shown to the right of the average profiles plot for one ORI SSR. B) As in A, but for SSRs not defined as origins (nonORI).

**Figure S10.**
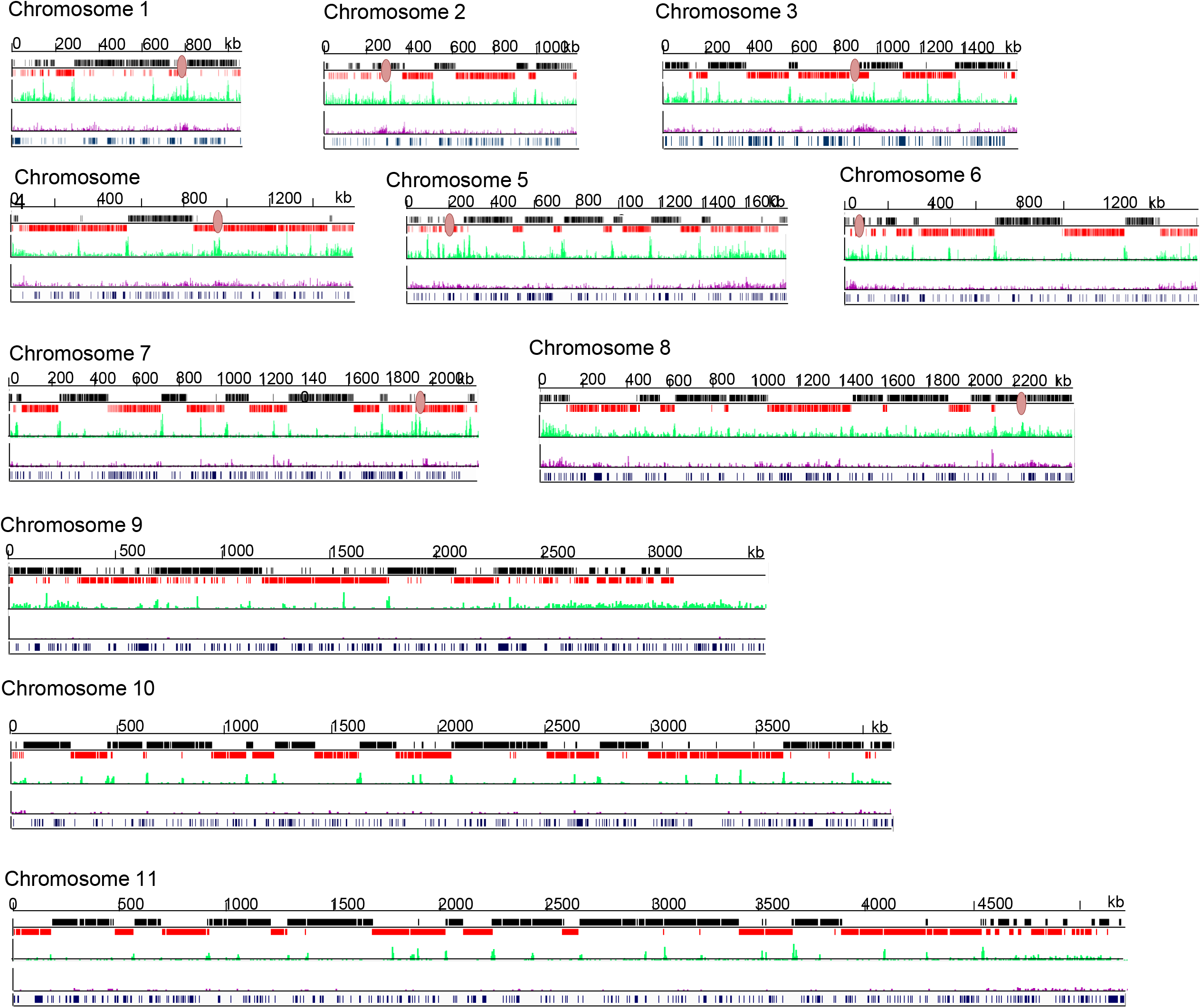
γH2A ChIP-seq signal is specifically enriched at sites of transcription initiation after TbRH2A depletion. Levels of γH2A ChIP-seq signal in TbRH2A RNAi induced sample coverage is plotted relative to uninduced sample coverage (each first normalised to input samples) for 24 hr (purple, low track) and 36 hr (green, upper track) time points in all 11 Mb-sized chromosomes. Scale: 1-3 fold-change. Other annotations are as in Fig. S4.

**Figure S11.**
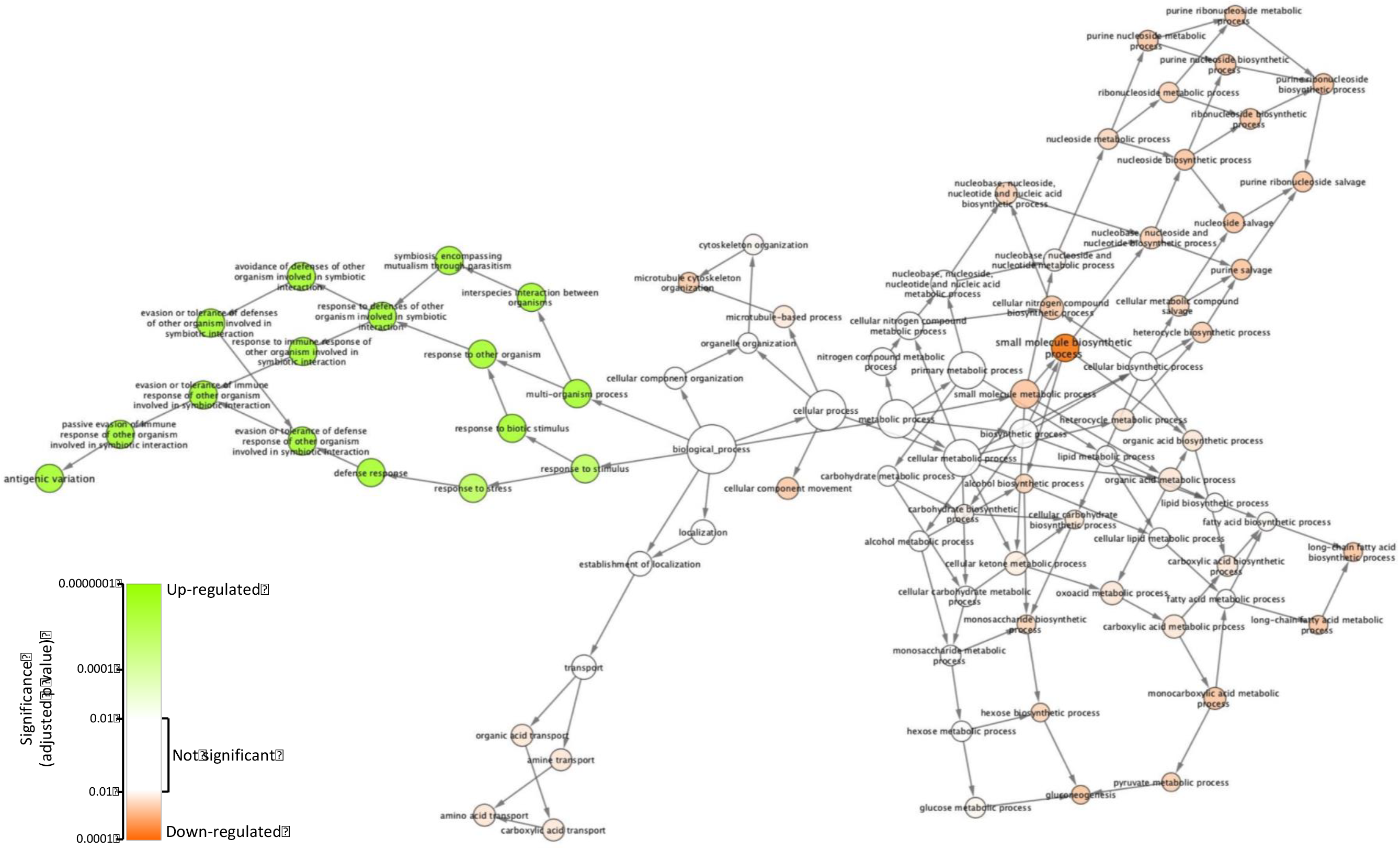
Gene ontology analysis reveals genes associated with antigenic variation are up-regulated after TbRH2A depletion, and those associated with small molecular biosynthesis pathways are down-regulated. Gene ontology analysis of biological process terms associated with genes found to be significantly up-regulated or down-regulated in RNA-seq analysis comparing RNA abundance after 36 hr TbRH2A of depletion via RNAi compared with uninduced cells. Parent terms are also depicted. Terms found to be enriched in the upregulated and downregulated genes are coloured green and orange, respectively. Colour intensity indicates significance, which was determined as adjusted p value. Terms with values > 0.01 were deemed not significant (white).

**Figure S12.**
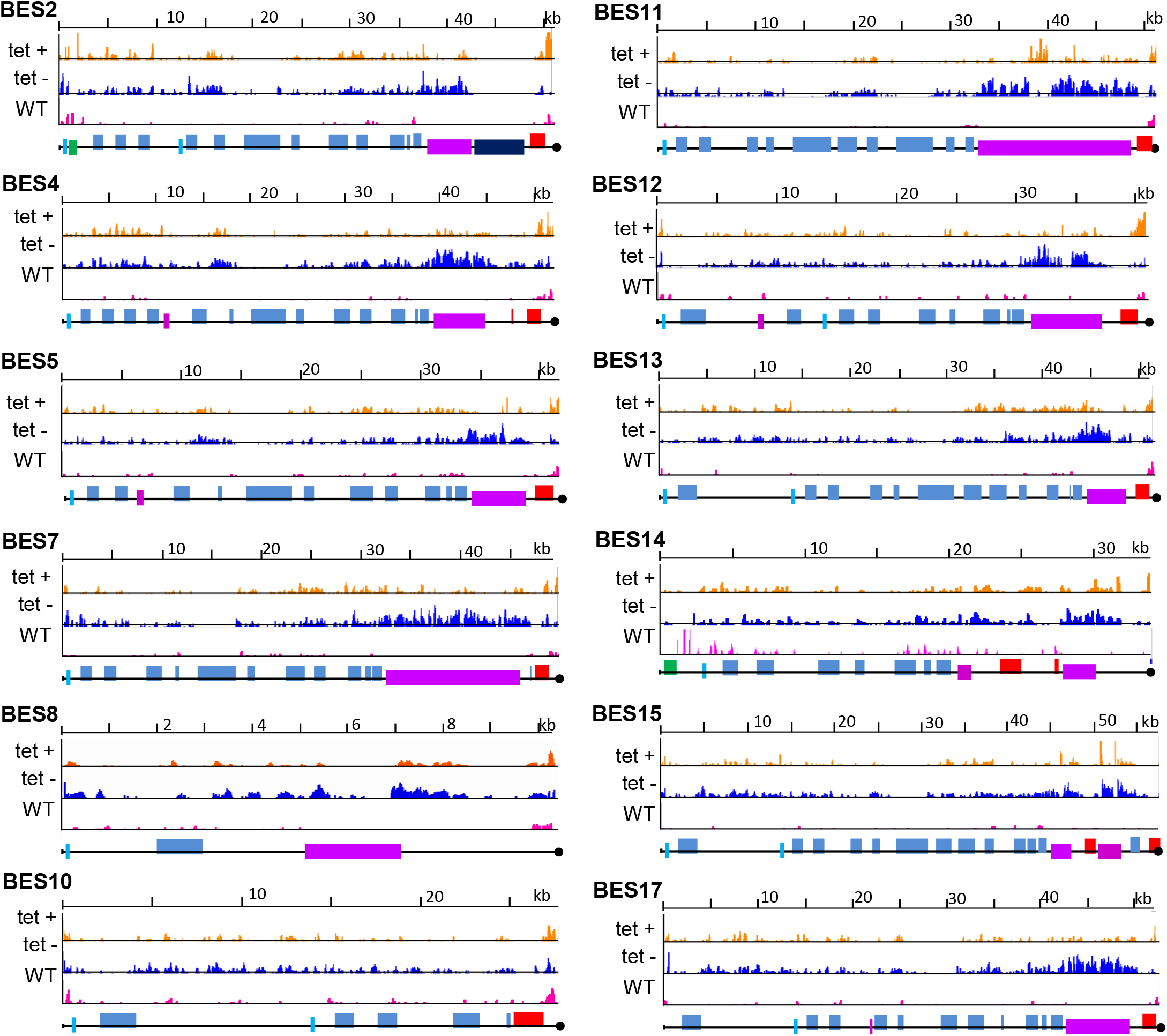
DRIP-seq signal is enriched across the BESs after TbRH2A depletion. DRIP-seq signal coverage (normalised to input coverage) is plotted across the BESs for WT (pink), TbRH2A uninduced (blue; tet -) and induced (orange, tet +) cells. The lowest track shows the structure of each BES; promoters (cyan), ESAGs (blue), 70-bp repeats (purple), VSGs (red) and other genes (green) are annotated as boxes. Black circles denote the end of the BES sequence assembly.

**Figure S13.**
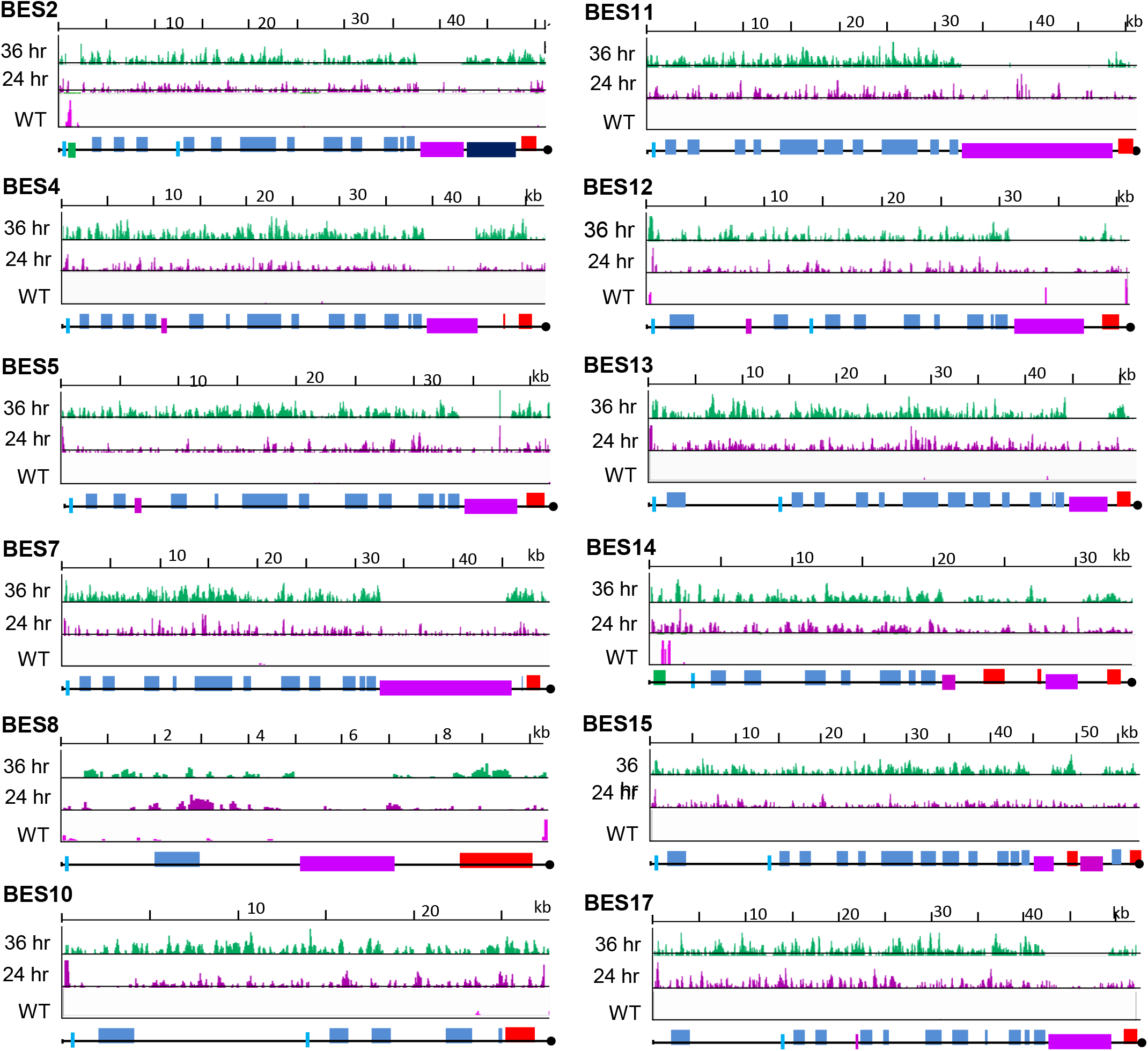
γH2A ChIP-seq is enriched across the BESs after TbRH2A depletion. Levels of γH2A ChIP-seq signal in TbRH2A RNAi induced sample coverage is plotted relative to uninduced sample coverage (each first normalised to input samples) for 24 hr (purple, low track) and 36 hr (green, upper track) time points, across each BESs. Scale, 1-3 fold-change. The lowest track shows the structure of each BES; promoters (cyan), ESAGs (blue), 70-bp repeats (purple), VSGs (red) and other genes (green) are annotated as boxes. Black circles denote the end of the BES sequence assembly.

**Figure S14.**
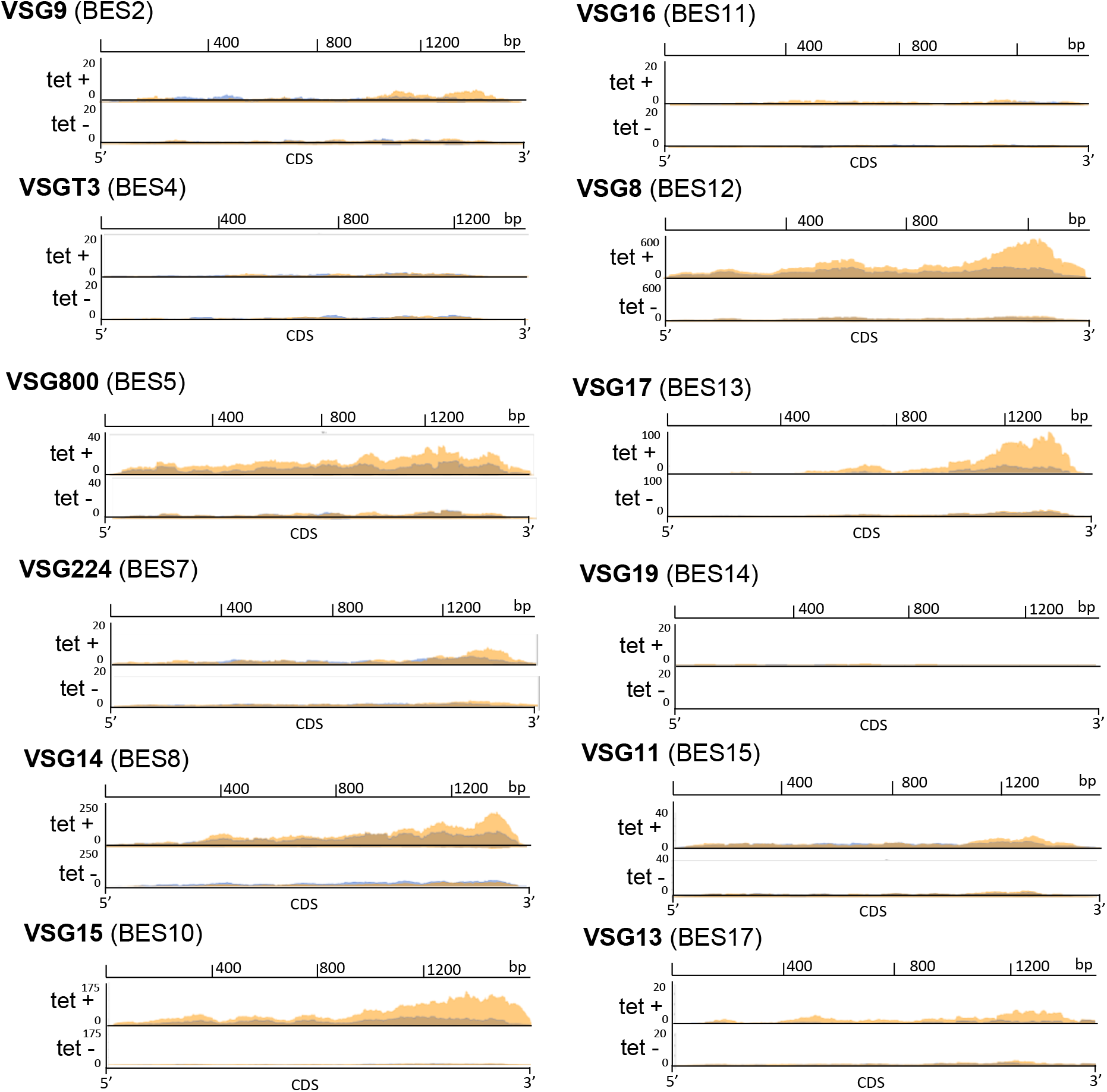
Silent BES-housed VSGs are transcribed after TbRH2A depletion. RNA-seq read coverage is plotted over the coding regions of VSGs housed within silent BESs for two independent replicates (blue and orange) of TbRH2A RNAi parasites grown for 36 hr in the absence (tet -) or presence (tet +) of tet-induction. The VSG names and BESs in which they are contained are indicated.

